# A general theory of individuated multicellularity

**DOI:** 10.1101/016840

**Authors:** Felipe A. Veloso

**Affiliations:** Center for Genomics and Bioinformatics, Faculty of Science, Universidad Mayor, Santiago, Chile

## Abstract

Changes in gene expression are thought to regulate the differentiation process intrinsically through complex epigenetic mechanisms. In fundamental terms, however, this assumed regulation refers only to the intricate propagation of changes in gene expression or else leads to logical inconsistencies. The evolution and self-regulatory dynamics of individuated multicellularity also lack a unified and falsifiable description. To fill this gap, I computationally analyzed publicly available high-throughput data of histone H3 post-translational modifications and mRNA abundance for different *Homo sapiens, Mus musculus,* and *Drosophila melanogaster* cell-type/developmental-periods samples. My analysis of genomic regions adjacent to transcription start sites generated a profile from pairwise partial correlations between histone modifications controlling for the respective mRNA levels for each cell-type/developmental-period dataset. I found that these profiles, while explicitly uncorrelated to transcript abundance by construction, associate strongly with cell differentiation states. This association is not expected if cell differentiation is, in effect, regulated by epigenetic mechanisms. Based on these results, I propose a falsifiable theory of individuated multicellularity, which relies on the synergistic coupling across the extracellular space of two stochastically independent “self-organizing” systems constraining histone modification states at the same sites. This theory describes how the multicellular individual—understood as an intrinsic, higher-order constraint—emerges from proliferating undifferentiated cells, and may explain the intrinsic regulation of gene transcriptional changes for cell differentiation and the evolution of individuated multicellular organisms.

---

Abbreviations: TSS, transcription start site; ChIP-seq, chromatin immunoprecipitation followed by high-throughput sequencing; RNA-seq, transcriptome high-throughput sequencing; FPKM, fragments per transcript kilobase per million fragments mapped.

## Introduction

Cell differentiation, if seen as a motion picture in fast-forward, intuitively appears to be a teleological or “end-directed” process, its *telos* or “end” being the multicellular organism in its mature form. The first step for a scientific explanation of this apparent property was given when Conrad H. Waddington proposed his epigenetic landscape model. Influenced by earlier developments in dynamical systems theory [1], Waddington’s model showed cell differentiation to be potentially predictable or at least explainable without any teleological reference [2].

The dynamics of the cell differentiation process have been associated with changes in chromatin states and concurrent heritable changes in gene expression that are uncorrelated to changes in the DNA sequence, and therefore defined as epigenetic changes [3, 4]. In some cases, these changes can be regulated extrinsically with respect to the developing organism, as observable in eusocial insects (e.g., a female honeybee larva develops into a worker or a queen depending on the royal jelly diet it is fed [5]). Yet most key changes in gene expression for cell differentiation are not only independent from, but are even robust with respect to extrinsic variables. This indicates that cell differentiation is fundamentally an intrinsically regulated process, for which no falsifiable theory has emerged from the epigenetic framework. Due to our lack of understanding of the precise regulatory dynamics, this process has also been dubbed “The X-files of chromatin” [6].

To unravel these X-files, we have to look critically at (i) the regulation of cell differentiation as it is understood today, (ii) the non-genetic information capacity of primordial cells (zygotes, spores, or buds), and (iii) what is assumed to be pre-specified developmental information content in those primordial cells. Modern science regards cell differentiation fundamentally as a dynamical system, where a fixed rule governs the transition between the realizable states of a complex network of molecular mechanisms. Ranging from low-order molecular interactions to chromatin higher-order structural changes [7, 8, 9, 10], these epigenetic mechanisms not only propagate changes in gene expression at different loci as cells proliferate but, importantly, are also hypothesized to regulate the cell differentiation process intrinsically. This hypothesis is accepted as a well-established fact (as illustrated in [11]) even though the epigenetic mechanisms involved in cell differentiation have not been fully elucidated. However, this epigenetic regulation hypothesis leads to severe explanatory limitations and may even entail logical inconsistencies.

If one assumes that this hypothesis is true in its strictest sense, one accepts that gene self-regulation is a teleological property of cell differentiation. For example, one might assume that a certain *gene A* is an explanatory component of the general self-regulatory property once a researcher who modifies the expression levels of *gene A* in a given organism elucidates how these changes activate or repress the expression of a specific *gene B, gene C,* and *gene D* during differentiation. However, this assertion overlooks that the researcher, not *gene A*, was the true regulator by purposefully imposing certain transcriptional states (on *gene A,* and by means of *gene A,* also *gene B, gene C,* and *gene D*). Yet, no extrinsic human regulator is needed for the natural process, which raises the question of what system is truly regulating *gene B, gene C, gene D* AND *gene A*—and by extension, all genes for cell differentiation. Moreover, accounting for the regulation of transcriptional states at a gene locus by previous transcriptional states at other gene loci—in the same cell or any other—is only a non-explanatory regress (Fig. S1).

If one assumes that the epigenetic regulation hypothesis is true in a loose sense, one has to use “self-regulation” only as a placeholder when referring to a certain class of molecular mechanisms propagating changes in gene expression. In this context, a proposed “epigenator”—a transient signal which probably originates in the environment of the cell—would trigger the epigenetic phenotype change after being transduced into the intracellular space [12]. However, if all “epigenators” in the developing organism are extrinsic to it, self-regulation is *ipso facto* unexplainable. Otherwise, the critical signaling property of an “epigenator” (i.e., what it refers to and how it does so) is left unexplained.

The question arises if it is possible that critical changes within a developing organism, and the intrinsic regulation of such changes, are completely different processes at the most fundamental level. Specifically, intrinsic regulation may not be a molecular mechanism correlating critical changes in gene expression within a developing organism but instead may involve particular *constraints* (understood as local thermodynamic boundary conditions that are level-of-scale specific) on those changes. Importantly, these particular constraints—imposed by the regulatory system we look for—are *stochastically independent* from the changes this system is supposed to regulate; otherwise the system is fundamentally just an additional mechanism propagating gene expression changes more or less extensively (depending, for example, on the presence of feedback loops) instead of *regulating* them. This explanatory limitation is inescapable: a nonlinear propagation of changes in gene expression only implies either a nonlinear dependence between those changes—describable by a dynamical systems model such as the epigenetic landscape—or chaotic behavior, not a *regulated* propagation of changes. Moreover, intrinsic regulation cannot be explained in terms of any mechanism, machine (e.g., autopoietic [13]), or any “self-organizing” system because all mechanisms, machines and “self-organizing” systems entail an explicit deterministic or stochastic dependence between all their component dynamics. Notably, however, the existence of “self-organizing” systems [14] is a necessary condition for the intrinsic regulatory system of cell differentiation under the theory I propose here.

Regardless of the explanatory limitations inherent to the epigenetic landscape, it is generally believed that most, if not all, non-genetic information for later cell differentiation is “hardwired” in the primordial cells. If these cells indeed contain all this information, including that for intrinsic regulation [15, 16], the previously discussed explanatory gap could, in principle, be filled. Asymmetric early cleavage, shown to be able to resolve a few cell lineage commitments (into six founder cells) in the nematode *Caenorhabditis elegans* [17], supports this possibility at first glance, but a closer look at the developmental complexity of this simple metazoan model organism suggests otherwise: the hermaphrodite *C. elegans* ontogeny yields 19 different cell types (excluding the germ line) in a total of 1,090 generated cells [18]. From these two parameters alone, the required information capacity for the entire process can be estimated to be at least 983bit (see details in Table S1). Furthermore, this is a great underestimation because uncertainty remains with respect to at least two other variables, namely space and time. Therefore, the non-genetic information capacity necessary for the entire cell differentiation process far exceeds the few bits shown to be accounted for by epigenetic mechanisms. On the other hand, extrinsic constraints (e.g., diet-dependent hierarchy determination in eusocial insects [5], temperature-dependent sex determination in reptiles [19], or maternal regulation of offspring development [20]) do not account for all developmental decisions. These considerations highlight that certain *intrinsic* constraints must be identified to account for all the necessary non-genetic information in terms of capacity, which is measurable in units such as bits and content, which must account for *how* each developmental decision is made.

The question also arises on how it is possible for an entire organism to develop from *any* totipotent cell, and for embryonic tissues to develop from *any* pluripotent stem cell, if the information for all cell fate decisions is contained in the primordial cell. The recently proposed “epigenetic disc” model for cell differentiation, under which the pluripotent state is only one among many metastable and directly interconvertible states [21], reflects the need to account for the significant dependence of developmental information on the cellular context.

Although David L. Nanney anticipated in 1958 that more important to development than the heritable material itself is the *process* by which heritable material may manifest different phenotypes [22], Waddington’s epigenetic landscape prevailed and ever since developmental biology has built upon the premise that primordial cells are indeed complete blueprints of the mature organism (allowing for some limited degree of stochasticity [23, 24] and extrinsic influence as described previously). Thus, the epigenetic landscape framework is not only fundamentally unable to explain the intrinsic regulatory dynamics of cell differentiation, but has also lead research to ignore or reject the necessary *emergence* of developmental information content during ontogeny.

To shed light into “The X-files of chromatin,” I designed and conducted a computational analysis of the combinatorial constraints on histone H3 post-translational modification states, referred to as histone H3 crosstalk, given their strong statistical relationship with transcriptional levels [25]. As the data source, I used publicly available tandem datasets of ChIP-seq on histone H3 modifications and RNA-seq on mRNA for *Homo sapiens, Mus musculus,* and *Drosophila melanogaster* cell-type, developmental-period, or developmental-time-point samples. The basis of the analysis was to define, from genomic coordinates adjacent to TSSs, a numeric profile *crosstalk that is non-epigenetic* (ctalk_non_epi), which is an *n*-tuple or ordered list of values with the sign and strength of partial correlations (i.e., normalized covariance) of histone H3 modification levels *once controlling for transcript abundance* (e.g., Cor(H3K27ac_-200,H3K4me1_-400|FPKM)). Thus, the ctalk_non_epi profile represents for any given sample and within TSS-adjacent genomic regions, the pairwise histone H3 crosstalk that is *stochastically independent* from (i.e., explicitly uncorrelated with) the transcriptional “identity” of that given cell type, developmental period, or developmental time point (Fig. 1).

**Fig. 1.**
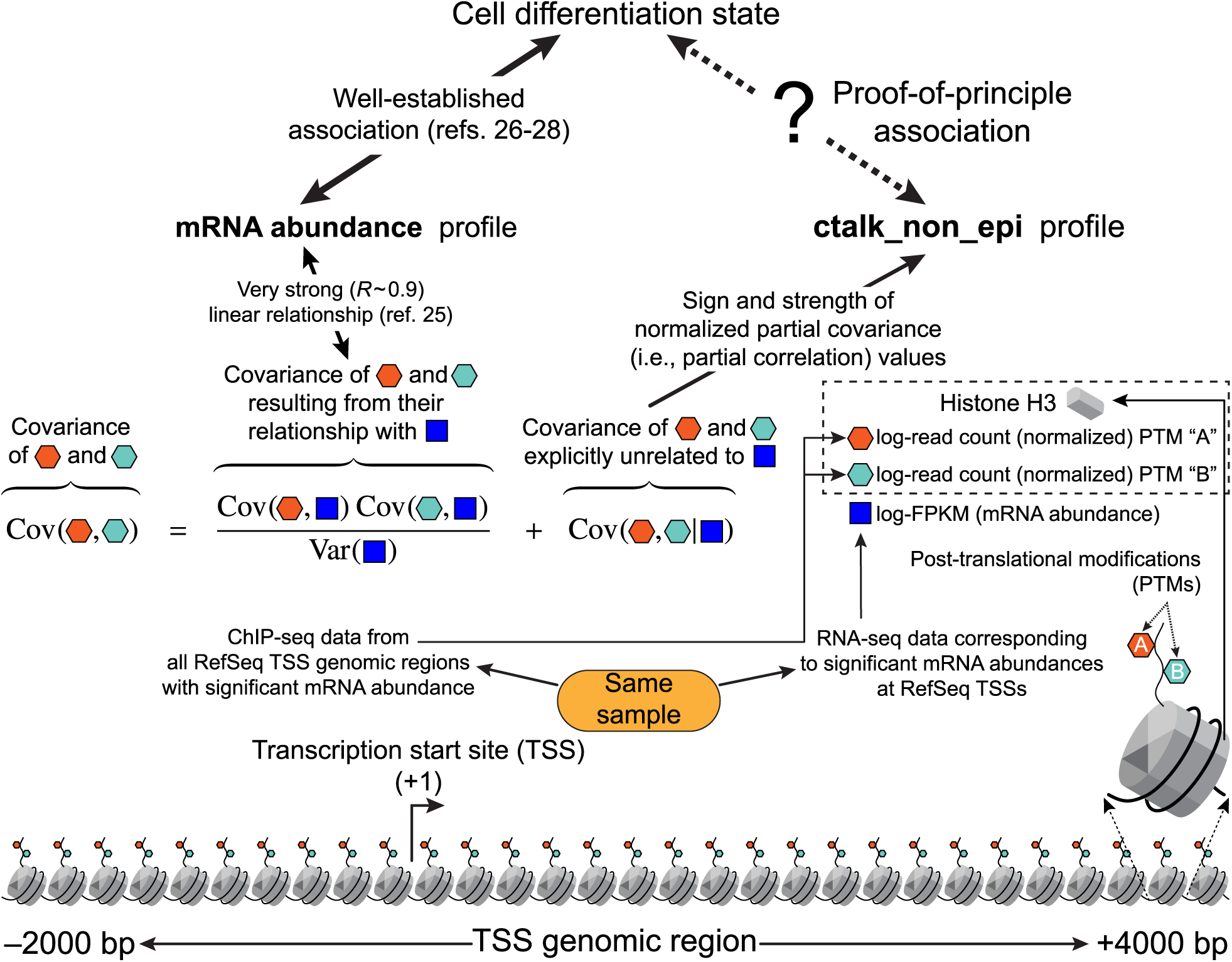
Scheme of the proof-of-principle hypothesis and the computational analysis for its testing. *ctalk_non_epi* profiles represent constraints on histone H3 crosstalk that are stochastically independent from mRNA levels. The association between these *ctalk_non_epi* profiles and cell-differentiation states established the proof of principle for the theory proposed in this paper.

## Results

Under the arguments presented in the introduction, the aim of my computational analysis was to test the following proof-of-principle, working hypothesis: *for a given cell differentiation state and within genomic regions adjacent to TSSs, the pairwise histone H3 crosstalk that is stochastically independent from transcriptional levels* (represented by the ctalk_non_epi profile) *associates with that differentiation state* (Fig. 1, black dashed arrow). Importantly, the null hypothesis (that is, no significant relationship exists between cell differentiation states and histone H3 crosstalk uncorrelated to mRNA levels) is further supported by the epigenetic landscape approach: if changes in mRNA levels not only associate with cell differentiation states [26, 27, 28] but also explain them completely, an additional non-epigenetic yet differentiation-associated type of constraint on histone H3 crosstalk is superfluous.

To test the proof-of-principle hypothesis, I applied hierarchical cluster analysis (HCA) on the ctalk_non_epi profiles for each organism analyzed. If there is no significant association between ctalk_non_epi profiles and cell differentiation states (i.e., if the null hypothesis is true), the obtained clusters should be statistically insignificant or else they should not associate with cell differentiation states. However, the results showed in all analyses performed that ctalk_non_epi profiles fell into statistically significant clusters that associate with cell differentiation states in *Homo sapiens*, *Mus musculus*, and *Drosophila melanogaster*. Moreover, ctalk_non_epi profiles associated with cell differentiation states at least as strongly as mRNA abundance profiles, whose obtained cluster structure was used for comparison and positive control. In sum, for all three organisms analyzed, the null hypothesis had to be consistently rejected in terms of a clear association between ctalk_non_epi profiles and cell differentiation states. This unambiguous result provides proof of principle for my theory, which requires differentiation-associated constraints on TSS-adjacent histone H3 crosstalk such that they are stochastically independent from mRNA levels.

### ctalk_non_epi profiles of embryonic stem cells differ significantly from those of differentiated cell types in human

Using data for nine different histone H3 modifications, I computed ctalk_non_epi profiles for six human cell types. From these, all profiles corresponding to differentiated cell types, namely HSMM (skeletal muscle myoblasts), HUVEC (umbilical vein endothelial cells), NHEK (epidermal keratinocytes), GM12878 (B-lymphoblastoids), and NHLF (lung fibroblasts) fell into the largest cluster. This cluster was also statistically significant as reflected in au (approximately unbiased) and bp (bootstrap probability) significance scores, which were greater than or equal to 95 (Fig. 2A, cluster #4), indicating that this cluster was also statistically significant. The ctalk_non_epi profile corresponding to H1-hESC (embryonic stem cells) was identified as the most dissimilar with respect to the other profiles, which are all differentiated cell types. These findings indicate that ctalk_non_epi profiles associate with cell differentiation states in *Homo sapiens*. Additionally, this association was clearer than the association observed between mRNA abundance profiles and cell differentiation states (Fig. S2A, cluster #3).

**Fig. 2.**
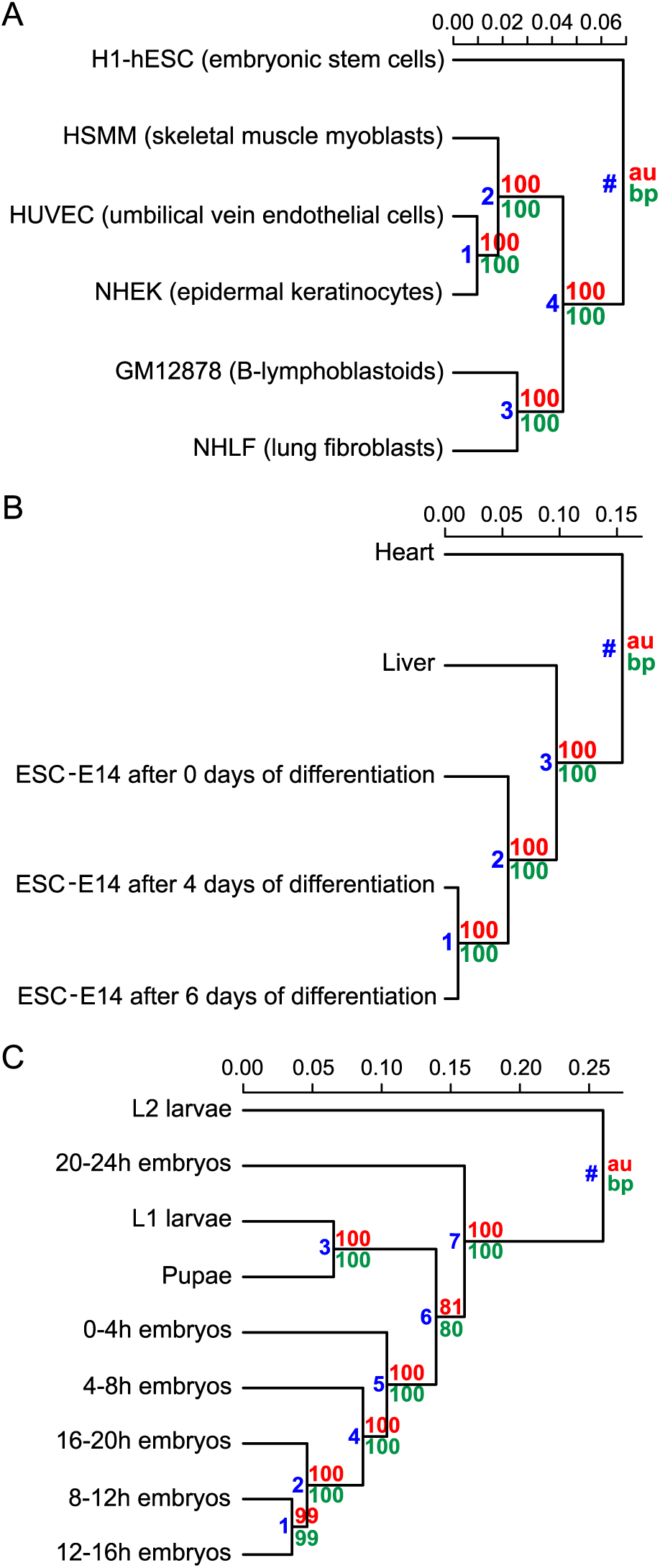
Hierarchical cluster analysis of *ctalk_non_epi* and mRNA abundance profiles. Organisms: *Homo sapiens* (A, B), *Mus musculus* (C, D), and *Drosophila melanogaster* (E, F). Metric: correlation (1 – *r*). Linkage method: UPGMA. Significance scores: au (approximately unbiased) and bp (bootstrap probability) [60]. Significant clusters were identified as those for which au and bp > 95. Cluster identification numbers are in blue.

### ctalk_non_epi profiles associate with cell differentiation states in mouse

The analysis for *Mus musculus* comprised five different histone H3 modifications in five cell types. The five cell type datasets analyzed were 8-weeks-adult heart, 8-weeks-adult liver, plus three datasets of E14 embryonic stem cells after zero, four, and six days of differentiation respectively. As in *Homo sapiens*, the ctalk_non_epi profiles for *Mus musculus* fell into significant clusters that associated with cell differentiation states. All three E14 ctalk_non_epi profiles were clustered into a significant, exclusive group (Fig. 2B, cluster #2) and within it, the profiles corresponding to latter time points (four and six days of differentiation) fell into another significant cluster (Fig. 2B, cluster #1). Additionally, the liver ctalk_non_epi profile was found to be more similar to the profiles of the least differentiated states than the heart profile (Fig. 2B, cluster #3). Overall, this analysis showed that the association between ctalk_non_epi profiles and cell differentiation states is also observable in *Mus musculus*.

### ctalk_non_epi profiles associate with developmental periods and time points in fruit fly

Similar to those from human and mouse data, ctalk_non_epi profiles were computed from data for six histone H3 modifications in nine periods/time points throughout *Drosophila melanogaster* development (0-4h, 4-8h, 8-12h, 12-16h, 16-20h and 20-24h embryos; L1 and L2 larval stages; pupae). As observed in human and mouse profiles, fruit fly ctalk_non_epi profiles fell into clusters that also associated strongly with the degree of cell differentiation. One significant cluster grouped ctalk_non_epi profiles of earlier developmental periods (Fig. 2C, cluster #5) apart from later development profiles. Two more significant clusters placed later time point ctalk_non_epi profiles (Fig. 2C, cluster #3) and separated the L2 larvae profile (Fig. 2C, cluster #7) from all other profiles. The mRNA abundance profiles in *D. melanogaster* yielded a general cluster structure that was much less consistent with developmental chronology than the ctalk_non_epi profiles (as exemplified by cluster #3 in Fig. S2C). Overall, these results indicate that the association between ctalk_non_epi profiles and cell differentiation states also holds in *Drosophila melanogaster* despite the limitations of the analysis imposed by the ChIP-seq/RNA-seq source data: unlike human and mouse data where each ctalk_non_epi profile represented a specific or almost specific differentiation state, each fruit fly dataset was obtained from whole specimens (embryos, larvae and pupae). Especially for later developmental stages, this implies that each ctalk_non_epi profile has to be computed from more than one partially differentiated cell type at the same developmental period.

## Discussion

Based on the proof of principle obtained, I propose a general theory of individuated multicellularity, which explains how gene expression is self-regulated for cell differentiation in extant multicellular lineages and how individuated multicellular lineages emerged throughout evolution. This theory describes how two constraint-generating or “self-organizing” systems elicit an emergent transition when they couple synergistically. The first system underpins the correlation between histone modification states and transcriptional states in the cell nucleus and the second system is a specific extracellular gradient generated by cell proliferation.

The theory is presented in thirteen parts that are described in terms of the evolution of an ancestor generic eukaryotic species *U* towards individuated multicellularity and in terms of the ontogenetic process starting from the primordial cell(s) of a generic individuated multicellular species *D*. Theoretical definitions and notation are listed below.

**Def. 1** **Extracellular space *S_E_***: the entire space in an organism or cell population that is not occupied by its cells at a given instant *t*. In the same logic, the following concepts must be understood in instantaneous terms.
**Def. 2** **Waddington’s constraints *C_W_***: the constraints associating elements of the spatially-specified molecular nuclear phenotype of the cell with the instantaneous transcription rates at the TSSs, provided these constraints are stochastically independent from changes in the genomic sequence.
**Def. 3** **Waddington’s embodiers *F_W_***: the elements of the spatially-specified molecular nuclear phenotype of the cell for which the Waddington’s constraints *C_W_* are significant (e.g., histone H3 post-translational modifications in TSS-adjacent genomic regions).
**Def. 4** **Nanney’s constraints *C_N_***: the constraints associating elements of the spatially-specified molecular nuclear phenotype of the cell with the Waddington’s embodiers *F*_W_, provided these constraints are stochastically independent from changes in the instantaneous transcription rates at TSSs (e.g., non-epigenetic histone H3 crosstalk in TSS-adjacent genomic regions, represented in the data analysis by the ctalk_non_epi profiles).
**Def. 5** **Nanney’s embodiers *F_N_***: the elements of the spatially-specified molecular nuclear phenotype of the cell for which the Nanney’s constraints *C_N_* are significant. Crucially, histone H3 post-translational modifications in the TSS-adjacent regions—as inferable from the Results—can be specified as Waddington’s embodiers *F_W_* and *also* as Nanney’s embodiers *F_N_*.
**Def. 6** **Nanney’s extracellular propagators** 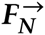: the elements of the entire molecular phenotype of the cell that are (i) secreted into the extracellular space *S_E_* and (ii) capable of eliciting a significant change (via facilitated diffusion and/or signal transduction) in Nanney’s constraints *C_N_* within other cells’ nuclei.
**Def. 7** **Gradient of Nanney’s extracellular propagators** 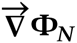: the vector whose components are the partial derivatives of the concentration Ф*_N_* of Nanney’s extracellular propagators 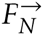 with respect to the spatial coordinates in *S_E_*.

### Part I: Before cell differentiation and its self-regulatory dynamics

When eukaryotic life was unicellular or organized into colonies at most, cell differentiation was not possible because the generation and propagation of changes in Nanney’s constraints *C_N_* (represented by the ctalk_non_epi profiles in the data analysis) were confined to each cell. In other words, Nanney’s extracellular propagators 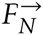 did *not* exist in the species *U* (Fig. S3A, top); a condition that also holds in extant unicellular or nonindividuated multicellular species. On the other hand, in primordial cells of the species *D* (as exemplified by zygotes, spores, or buds) Nanney’s extracellular propagators 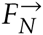 *do* exist, but self-regulated cell differentiation is not observed *yet* (Fig. S3A, bottom).

### Part II: Necessary alleles

At some point during evolution, certain *U* cell suffers a genetic change (Fig. S3A to S3B) such that this cell now features Nanney’s extracellular propagators 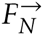. This genetic change implies that its genome now codes for all gene products necessary for the synthesis, facilitated diffusion, and/or signal transduction of the novel 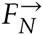. On the other hand, this novel genome is similar to the genome of any cell of the species *D* at any instant in that both genomes display the necessary alleles for 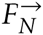. Importantly, these alleles—unprecedented for the species *U* and preexisting for the species *D*—are a *necessary but not sufficient* condition for individuated multicellularity.

### Part III: Diffusion flux of Nanney’s extracellular propagators and the geometry of the extracellular space

When the numbers of cells in the offspring of the ancestor *U* cell and of the primordial *D* cell(s) is small enough, diffusion flux of Nanney’s extracellular propagators 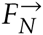 is fast enough to overtake the spatial constraints imposed by the relatively simple geometry of *S_E_*. As a consequence, the concentration of 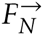 is relatively uniform in *S_E_*.

### Part IV: The emergent transition to individuated multicellularity and to cell differentiation

At some later instant, cell proliferation or embryonic growth yields a significantly large cell population for which diffusion flux of Nanney’s extracellular propagators 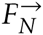 is no longer able to overtake the increasing spatial constraints in the extracellular space *S_E_*. Under these conditions a significant gradient 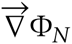 forms, in magnitude equal to or greater than the critical values *V_m_* (Fig. 3, bottom-left) or *V_d_* (Fig. 3, bottom-right)—anywhere in *S_E_*. As a consequence, Nanney’s extracellular propagators 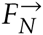 diffuse differentially into each cell, yielding unprecedented differential changes in Nanney’s constraints *C_N_* in the cells’ nuclei, not because of any cell, gene product or transcriptional profile, but because of the constraints imposed by the entire proliferating cell population on the diffusion flux of 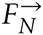 in *S_E_*. Consequently, because Nanney’s constraints *C_N_* are defined with respect to Waddington’s embodiers *F_W_* (e.g., histone post-translational modifications in TSS-adjacent genomic regions; see Fig. 3, center), the instantaneous transcription rates at TSSs change *differentially* and *irrespectively* of how gene transcriptional changes were *propagating* up to that instant and thus cells differentiate. In other words, instantaneous transcription rates will change by virtue of *true regulatory* constraints. Importantly, an extracellular gradient 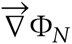of significant magnitude implies the availability of non-equilibrium Gibbs free energy which, as will be described later, is in fact partially used as *work* in the emergence of unprecedented information *content*, which is critical for multicellular individuation.

**Fig. 3.**
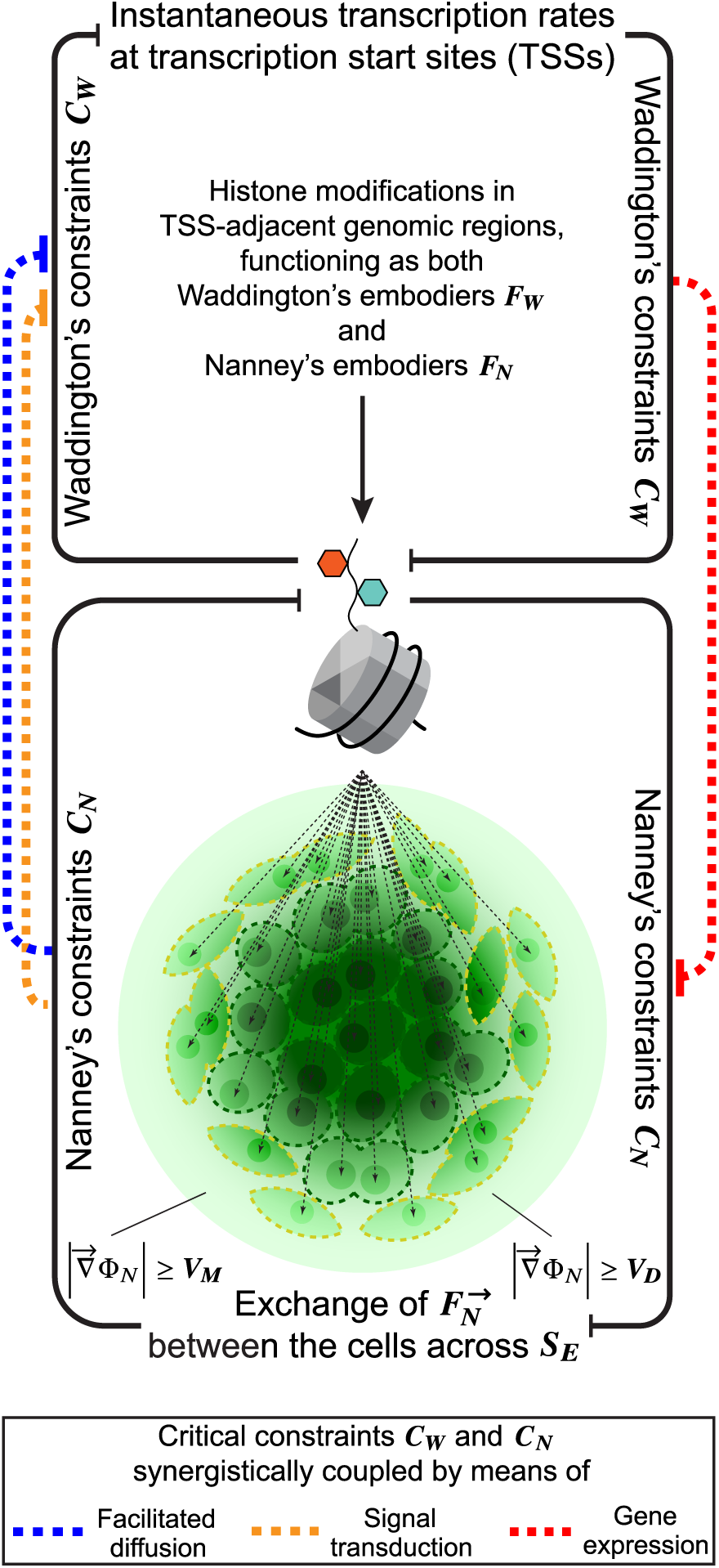
Emergent transition to individuated multicellularity and to cell differentiation. Intrinsic higher-order constraint emerges when significant gradients 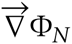 couple the lower-order and stochastically independent Waddington’s constraints *C_W_* and Nanney’s constraints *C_N_* synergistically across *S_E_*. Histone modification states in the TSS-adjacent genomic regions become thus constrained differentially by two stochastically independent “self-organizing” systems. This is how individuated multicellular lineages emerged throughout evolution and how cell differentiation emerges during ontogeny, all displaying a truly *self-regulated* dynamical regime.

### Part V: The true intrinsic regulator of cell differentiation

Contrary to what could be derived from Turing’s hypothesis [29], the theory hereby proposed does *not* regard the significant proliferation-generated extracellular gradient 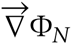 as the fundamental intrinsic regulator of the cell differentiation process. Whereas differential Nanney’s constraints *C_N_* are *regulatory constraints* with respect to Waddington’s constraints *C_W_* as described in Part IV (see Fig. 3, blue/orange dashed lines), the reciprocal proposition is also true. Namely, Waddington’s constraints *C_W_* are *stochastically independent* from Nanney’s constraints *C_N_*, thus *C_W_* are in turn *regulatory constraints* with respect to *C_N_*, e.g., by modifying the expression of protein channels, carriers, membrane receptors, or intracellular transducers necessary for the facilitated diffusion/signal transduction of Nanney’s extracellular propagators 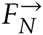 into the cells (Fig. 3, red dashed lines). Such reciprocity or synergy between two stochastically independent “self-organizing” systems coupled across *S_E_* by virtue of Nanney’s extracellular propagators 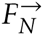 is what explains cell differentiation as a self-regulated process, not epigenetic mechanisms, which should be regarded as necessary propagators (yet not true regulators) of changes in gene expression. *Consequently, only if the stochastically independent Waddington’s constraints C_W_ and Nanney’s constraints C_N_ become synergistically coupled across the extracellular space S_E_, true intrinsic regulation on the cell differentiation process is possible* (Fig. 3). This corollary implies in turn that both histone modification states in TSS-adjacent genomic regions and transcriptional states are reciprocally cause and effect with respect to each other, thus providing also a plausible fundamental account of the more general and so far unresolved causal relationship between nuclear organization and gene expression (discussed in [30]). *The true intrinsic regulator of the cell differentiation process is the developing multicellular individual itself*. This is because the multicellular individual *is* the intrinsic (with respect to the cell population) and causally-efficacious higher-order *constraint* emergent from and regulating *ipso facto* Waddington’s constraints *C_W_* and Nanney’s constraints *C_N_* in what would be otherwise an arbitrarily complex colony of eukaryotic cells.

### Part VI: The major evolutionary breakthrough

Although necessary, the novel genetic sequences featured by individuated multicellular lineages were not the major evolutionary breakthrough. What was fundamentally unprecedented for life on Earth was the dynamical regime emerging for the first time, as described previously, in more than one lineage throughout evolution since extant individuated multicellular species constitute a paraphyletic group [31, 32]. The system displaying this novel dynamical regime is not a network of molecular mechanisms—however complex. Instead, it is an *intrinsic higher-order constraint* on component cells or multicellular *self*, emerging from the synergistic coupling of the lower-order Waddington’s constraints *Cw* and Nanney’s constraints *C_N_* across *S_E_* (Fig. 3, black lines). Because it is an intrinsic, higher-order *constraint,* this multicellular self or individual *can be causally-efficacious* when regulating its intrinsic dynamics or modifying its surroundings. The multicellular self described here is in turn a particular example of the generic *teleodynamic system* (a scientifically-tenable, causally-efficacious self or individual), proposed by Terrence W. Deacon in his *emergent dynamics* theory [33], which emerges when lower-order “self-organizing” systems become synergistically coupled (i.e., when the local thermodynamic boundary conditions required by each such system are generated by the others) and *then* is subject to and shaped by the interplay between its emergent properties and neo-Darwinian mechanisms (interplay henceforth referred to simply as evolution).

### Part VII: Unprecedented multicellular regimes

Once the necessary alleles for individuated multicellularity are present in some eukaryotic lineages (see Part II), further variation due to phenomena like mutation, gene duplication or alternative splicing make possible the emergence of a plethora of novel (teleodynamic) multicellular regimes and their respective developmental trajectories. Since a higher-order constraint is taking over the regulation of many changes in gene expression within individual cells, it is predictable that said cells lose some cell-intrinsic systems that were critical at a time when eukaryotic life was only unicellular, even when compared to their prokaryotic counterparts. In this context a result obtained over a decade ago acquires relevance: in a genome-wide study it was found that that the number of transcription factor genes increases as a power law of the total number of protein coding genes, with an exponent greater than 1 [34]. In other words, the need for transcription-factor genetic information increases faster than the total amount of genes or gene products it is involved in regulating. Intriguingly, the eukaryotes analyzed were the group with the smallest power-law exponent. This means that the most complex organisms require proportionally *less* transcription-factor information. With data available today [35], a reproduction I conducted of the aforementioned analysis allowed a robust confirmation: the power-law exponent for unicellular and nonindividuated multicellular eukaryotes is 1.33 ± 0.31 (based on 37 genomes). For individuated multicellular eukaryotes is 1.11 ± 0.18 (67 genomes). The loss of lower-order, cell-intrinsic regulatory systems in individuated multicellular organisms described above—in turn accounted for by the emergence of higher-order information content (see Part XI)—explains these otherwise counterintuitive differences in power-law exponents.

### Part VIII: What does ontogeny recapitulate?

As the key to the evolution of any multicellular lineage displaying self-regulated changes in gene expression for cell differentiation, the proposed theory holds the emergent transition, spontaneous from cell proliferation shortly after Nanney’s extracellular propagators 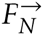 began to be synthesized and exchanged/transduced differentially through the cells’ membrane. Therefore, the theoretical description presented here rejects the hypothesis that any eukaryotic lineage displaying self-regulated cell differentiation evolved from gradual specialization and division of labor in single-cell colonies or aggregations [36, 37, 38, 32, 39, 40, 41, 42]. This rejection, however, does not imply that precedent traits (e.g., cell-cell adhesion) were unimportant for the later fitness of individuated multicellular organisms. Haeckel’s famous assertion is not rejected completely because it contains some truth: in every extant multicellular lineage, the self-sufficient, self-repairing, self-replicating, and self-regulating system described in this theory emerges over and over again from proliferating undifferentiated cells and presents itself to evolution ever since its phylogenetic debut. Therefore, at least in this single yet most fundamental sense, ontogeny does recapitulate phylogeny.

### Part IX: The role of epigenetic changes

Contrary to what the epigenetic landscape framework entails, under this theory the heritable changes in gene expression do not define, let alone explain by themselves, the intrinsic regulation of cell differentiation. The robustness, heritability, and number of cell divisions that any epigenetic change comprises are instead *adaptations* of the intrinsic higher-order constraint emergent from proliferating undifferentiated cells (i.e., the multicellular individual or self). These adaptations have been shaped by evolution after the emergence of each extant multicellular lineage and are in turn reproduced, eliminated, or replaced by novel adaptations in every successful ontogenetic process.

### Part X: Novel cell types, tissues and organs evolve and develop

Further genetic variation (e.g., mutation, gene duplication, alternative splice sites) in the novel alleles of the ancestor *U* cell or in the preexisting alleles of the primordial *D* cells imply than one or more 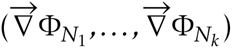 gradients form in *S_E_* with cell proliferation. A cell type *T_j_* will develop then in a subregion 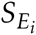 of the extracellular space *S_E_* when a relative uniformity of Nanney’s extracellular propagators is reached, i.e., when the gradients 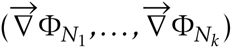 remain in magnitude under certain critical values 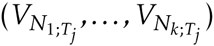 (see a two-cell-type and two-gradient depiction in Fig. S4). The constraint synergy underpinning this process can be exemplified as follows: gradients 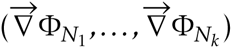 can elicit changes in gene expression in a number of cells, which in turn may promote the dissipation of the gradients (e.g., by generating a membrane surrounding the cells that reduces dramatically the effective *S_E_* size) or may limit further propagation of those gradients from *S_E_* into the cells (e.g., by repressing the expression of genes involved in the facilitated diffusion/signal transduction of 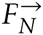 from *S_E_*). Thus, under this theory, cell types, tissues, and organs evolved sequentially as “blobs” of relative 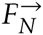 uniformity in regions 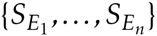 (i.e., regions of relatively small 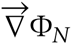 magnitude; see Fig. S5A) within *S_E_* displaying no particular shape or function—apart from being compatible with the multicellular individual’s survival and reproduction—by virtue of genetic variation (involved in the embodiment and propagation of Nanney’s constraints *C_N_*) followed by cell proliferation. Then, these 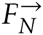-uniformity “blobs” were shaped by evolution from their initially random physiological and structural properties to specialized cell types, tissues, and organs (Fig. S5, gray dashed arrows) that *serve* the emergent intrinsic higher-order constraint proposed here as being the multicellular individual itself.

### Part XI: Emergent hologenic information and developmental robustness

A significant amount of information content has to *emerge* to account for robust and reproducible cell fate decisions and for the self-regulated dynamics of cell differentiation in general. Under this theory, such information content emerges when significant gradient or gradients 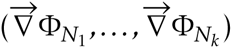 form at some point from proliferating undifferentiated cells, coupling synergistically Waddington’s constraints *C_W_* and Nanney’s constraints *C_N_* across *S_E_*. Crucially, this information is *not* genetic *nor* epigenetic. Instead, it is *about the multicellular individual as a whole*, understood as the emergent higher-order intrinsic constraint described previously, and also about the environmental constraints under which this multicellular individual develops. For this reason, I propose to call this emergent information *hologenic*—the suffix -genic may denote “producing” or “produced by”, which are both true under this theory. This is because the multicellular individual is, at each instant, not only interpreting hologenic information—by constraining its development into specific trajectories since its emergence given the current 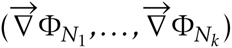 gradients in *S_E_* —but also actively generating novel hologenic information, e.g., when its very growth and the morphological changes in its differentiating cells modify the spatial constraints in *S_E_* and, as a consequence, the 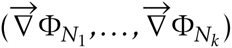 gradients. This causal circularity is similar to that described in Part V because it is underpinned by the same logic of constraint reciprocity (i.e., a teleodynamic relationship [33, 43]). Thus, in the most fundamental sense, *cell differentiation is an interpretive process*, not the replication or inheritance of any contentless molecular “code”, as David L. Nanney had indirectly anticipated [22]; its *defining interpreter of information*—endogenous such as hologenic information or exogenous such as that in royal jelly feeding [5]—*is the developing individual itself*. The *content* of hologenic information must be understood as *the otherwise realizable states that become constrained in the dynamics of the multicellular individual by the synergistic coupling of Waddington’s constraints C_W_ and Nanney’s constraints C_N_ across S_E_*, not the interactions of molecular substrates conveying such information (by embodying and propagating the critical constraints). Additionally, since the gradients 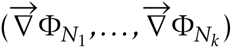 conveying hologenic information depend on no cell in particular but on the spatial constraints imposed by the entire cell population or embryo, cell differentiation can be robust with respect to moderate perturbations such as some cell loss.

### Part XII: Ontogeny ends and cell differentiation “terminates”

If cell differentiation emerges with the proliferation of (at the beginning, undifferentiated) cells, why should it terminate for any differentiation lineage? What is this “termination” in fundamental terms? These are no trivial questions. As an answer to the first, zero net proliferation begs the fundamental question; to the second, a “fully differentiated” condition fails to explain the existence of adult stem cells. Under this theory, cell differentiation “terminates” in any given region 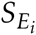 of the extracellular space if a stable or metastable equilibrium is reached where at least one of the two following conditions is true:

(a) The critical gradients 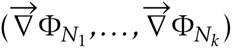 remain in magnitude under certain values 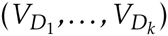 in 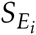 (Fig. S5B, left).

Condition (a) can be reached for example when development significantly changes the morphology of the cells by increasing their surface-to-volume ratio, because such increase removes spatial constraints in 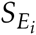 that facilitate the formation/maintenance of the gradients. Thus, under this theory, one can predict a *significant positive correlation between the degree of differentiation of cells and their surface-to-volume ratio* and also a *significant negative correlation between cell potency/regenerative capacity and that ratio*, once controlling for cell characteristic length. Condition (a) can also be reached when the current—since they are instantaneously defined—Nanney’s extracellular propagators 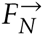 are non-functional or not expressed.

(b) The critical gradients 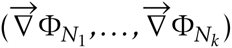 are significant but unable to modify Nanney’s constraints *C_N_* differentially in the cells’ nuclei (Fig. S5B, right).

Condition (b) can be reached when the cell differentiation process represses at some point the expression of the protein channels, carriers, membrane receptors or intracellular transducers necessary for the facilitated diffusion/signal transduction of the *current* Nanney’s extracellular propagators 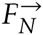, i.e., the cells become “blind” to the significant gradients 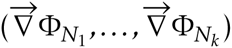 if they exist. Importantly, the stability of these equilibrium conditions will depend on the cells’ currently expressed phenotype. For example, an adult multipotent or pluripotent stem cell may differentiate if needed [44] or some differentiated cells may dedifferentiate given certain stimuli [45] (metastable equilibrium), whereas a fully differentiated neuron does not (very stable equilibrium). These examples underscore in turn that the *telos* of cell differentiation is not a “fully differentiated” state but, as this theory explains, the instantaneous, intrinsic higher-constraint, which is the multicellular individual as a whole.

### Part XIII: The evolutionarily-shaped multicellular telos

Under this theory, ontogeny is an emergent, evolutionarily-shaped and instantaneously-defined (i.e., logically consistent) teleological process. The reason why it intuitively (yet inconsistently in terms of causal logic) appears to be “directed” to and by the organism’s mature form is that the intrinsic higher-order constraint—the true (instantaneous) *telos* described previously—and the hologenic information content emerging along with it, are exerting *efficacious causal power* (i.e., agency) on the ontogenetic process. Although critical lower-order processes, such as the propagation of changes in gene expression, are decomposable into molecular interactions, the “end-directed” causal power of self-regulation in cell differentiation is *not* because its *telos* or “end” (Fig. S1, top right) is an intrinsic higher-order *constraint* or *local and level-of-scale-specific thermodynamic boundary condition* (previously described by T.W. Deacon also as a *“dynamical equivalent of zero*” [33]). Evolution has shaped the content of hologenic information (from the initial “blobs” of relative 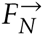 uniformity in *S_E_*, see Fig. S5A) by capturing the lower-order genetic constraints it is ultimately emergent from, not any particular molecules as media for their embodiment, media that should be regarded in this context as *means* to the multicellular *telos*. This explanation also implies a trade-off between cell independence and cell phenotypic complexity/diversity: the multicellular *telos* offloads regulatory work the cells were performing individually (as described in Part VII), allowing them to use that free energy surplus for sustaining more complex and diverse phenotypes but also making them more dependent on the multicellular *telos* they *serve.* The description for the evolution of cell types, tissues and organs based on initial “blobs” of relative 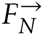 uniformity in *S_E_* together with the predicted positive correlation between degree of cell differentiation and cell surface-to-volume ratio suggest an additional and more specific evolutionary implication: the high surface-to-volume ratio morphology needed for neuron function was only to be expected as a trait of highly differentiated cells in the evolution of multicellularity, provided no high-fitness traits impede the tinkering with substantial increases of the cells’ surface-to-volume ratio. Together with the predicted negative correlation between cell potency and cell surface-to-volume ratio, this caveat suggests that if a multicellular lineage is constrained to display low cell surface-to-volume ratios, cell potency and regenerative capacity will be higher. These multicellular lineages can thus be expected to have a comparatively lower complexity but higher cell potency and robustness to extrinsic damage, as seen in the plant lineage (bound to exhibit low cell surface-to-volume ratios by virtue of rigid and concave cell walls).

The synergy in the coupling of Waddington’s constraints *C_W_* and Nanney’s constraints *C_N_* across *S_E_* described in this theory does not preclude that cell differentiation may display phases dominated by proliferation and others dominated by differentiation itself: whereas significant gradients 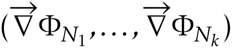 form in *S_E_* at some point given enough cell proliferation, it is also true that the exchange of Nanney’s extracellular propagators 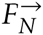 across *S_E_* is constrained by the dynamics of facilitated diffusion and/or ligand-receptor binding, which are saturable. Any representative simulation of cell differentiation according to this theory, however simple, will depend on an accurate modeling of the lower-order constraints it emerges from. The proposed theory also encompasses coenocytic or “syncytial” stages of development, where cell nuclei divide in absence of cytokinesis, as observed in *Drosophila*. In such stages, 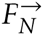 have to be operationally redefined as Nanney’s *extranuclear* propagators, while still maintaining their fundamental defining property: the ability to elicit changes in Nanney’s constraints *C_N_* within other nuclei. Related to this theory, evidence has already been found for tissue migration across a migration-generated chemokine gradient in zebrafish [46, 47].

### Falsifiability

Popper’s criterion of falsifiability will be met by providing three experimentally-testable predictions, which follow directly from the presented theory:

1. If undifferentiated stem cells or their differentiating offspring are extracted continuously from a developing embryo at the same rate they are proliferating, the undifferentiated cells will not differentiate or the once differentiating cells will enter an artificially-induced diapause or developmental arrest. A proper experimental control will be needed for the effect of the cell extraction technique itself, in terms of applying the technique to the embryo but extracting no cells.
2. A significant positive correlation will be observed between the overall cell-type-wise dissimilarity of Nanney’s constraints *C_N_* in an embryo (representable by ctalk_non_epi profiles) and developmental time.
3. If any molecule *M* is (i) specifiable as a Nanney’s extracellular propagator 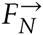 during a certain time interval for certain cells of a individuated multicellular species *D* and (ii) also synthesized by a unicellular (or nonindividuated multicellular) eukaryote species *U*, then experiments will fail to specify *M* as a Nanney’s extracellular propagator 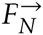 for the species *U*.

The strongest verifiable—yet not falsifiable—prediction that follows from the theory is *the existence of Nanney’s extracellular propagators 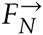 in any individuated multicellular species.* Since these propagators are instantaneously defined, their identification should be in the form “molecule *M* is specifiable as a Nanney’s extracellular propagator 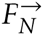 of the species *D* in the cell, cell population, or cell type *T_j_* at the developmental time point *t* (or the differentiation state *s*)”. This will be verified if, for instance, an experiment shows that the ctalk_non_epi profile (which represents Nanney’s constraints *C_N_*) of these *T_j_* cell or cells varies significantly when exposed to differential concentrations of *M* in the extracellular medium. Importantly, there is no principle in this theory precluding a molecule *M* that is secreted into the extracellular space *S_E_* and that activates or represses the expression of certain genes in other cells from being also specifiable as a Nanney’s extracellular propagator 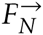. In other words, more likely than the discovery of previously undescribed molecules will be *the verification of the ability of some known secreted molecules to elicit changes in Nanney’s constraints C_N_ within other cells’ nuclei.*

### Corollaries

Hypotheses and predictions (not involving falsifiability) that can be derived from the proposed theory—regarding topics such as the evolution and development of the extracellular matrix (ECM), natural developmental arrests or diapauses, the regenerative response to wounds, the effects of microgravity on development, and the onset of cancer—can be found in *SI Corollaries.*

### Concluding remarks

Here, I show that scientifically tenable teleology in nature can emerge only from local and level-of-scale-specific thermodynamic boundary conditions (i.e., constraints) that are *stochastically independent* with respect to each other at certain critical sites such as those for histone post-translational modifications in TSS-adjacent genomic regions. The only way such requisite of stochastic independence can be fulfilled intrinsically is if a higher-order constraint emerges from the synergistic coupling of lower-order constraint generating systems, an emergent transition first described by T.W. Deacon. Whereas this thermodynamically spontaneous, intrinsic constraint—the logically-consistent *telos*—is dependent on molecular substrates embodying it at any instant, these substrates can be added, replaced or even dispensed with at any instant as long as the *telos* is preserved. For all these reasons, the individuated multicellular organism described in this theory is no mechanism, machine, or “self-organizing” system of any type as such systems entail an *explicit deterministic or stochastic dependence* between their component dynamics. Thus, the emergence of individuated multicellularity throughout evolution and in every successful ontogenetic process has been—and still is—the emergence of unprecedented higher-order intrinsic constraints or *selves* in the natural world; *selves* whom no mechanism, machine, or “self-organizing” system could ever be.

## Materials and methods

### Data collection

The genomic coordinates of all annotated RefSeq TSSs for the hg19 (human), mm9 (mouse), and dm3 (fruit fly) assemblies were downloaded from the UCSC (University of California, Santa Cruz) database [48]. Publicly available tandem datafiles of ChIP-seq (comprising 1×36 bp, 1×50 bp, and 1×75 bp reads, depending on the data series) on histone H3 modifications and RNA-seq (comprising 1×36 bp, 1×100 bp, and 2×75 bp reads, depending on the data series) for each analyzed cell sample in each species were downloaded from the ENCODE, modENCODE or the SRA (Sequence Read Archives) database of the National Center for Biotechnology Information [49, 50, 51, 52, 53, 54, 55].

### ChIP-seq read profiles and normalization

The first steps in the EFilter algorithm by Kumar *et al*.—which predicts mRNA levels in log-FPKM with high accuracy [25]—were used to generate ChIP-seq read signal profiles for the histone H3 modifications data.

### ChIP-seq read count processing

When the original format was SRA, each datafile was pre-processed with standard tools fastq-dump, bwa aln, bwa samse, samtools view, samtools sort and samtools index to generate its associated BAM (Binary Sequence Alignment/Map) and BAI (BAM Index) files. Otherwise, the tool bedtools multicov was applied (excluding failed-QC reads and duplicate reads by default) directly on the original BAM file (the BAI file is required implicitly) to generate the corresponding read count file in BED (Browser Extensible Data) format.

### RNA-seq data processing

The processed data were mRNA abundances in FPKM at RefSeq TSSs. When the original file format was GTF (Gene transfer Format) containing already FPKM values (as in the selected ENCODE RNA-seq datafiles for human), those values were used directly in the analysis. When the original format was SAM (Sequence Alignment/Map), each datafile was pre-processed by first sorting it to generate then a BAM file using samtools view. If otherwise the original format was BAM, mRNA levels at RefSeq TSSs were then calculated with FPKM as unit using the Cufflinks tool [56].

### Computation of ctalk_non_epi profiles

If the variables *X_i_* (representing the signal for histone H3 modification *X* in the genomic bin *i* ∈ {“−2000”,..., “3800”}), *Y_j_* (representing the signal for histone H3 modification *Y* in the genomic bin *j* ∈ {“−2000”,..., “3800”}) and *Z* (representing log_2_-transformed FPKM values) are random variables, then the covariance of *X_i_* and *Y_j_* can be decomposed in terms of their linear relationship with *Z* as follows:

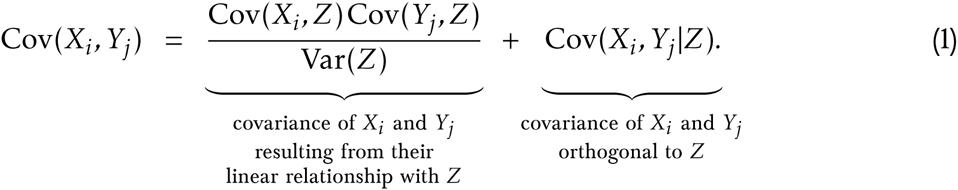

After log-transforming the *X_i_*, *Y_j_* and *Z* data, each pairwise combination of bin-specific histone H3 modifications {*X_i_, Y_j_*} contributed to the ctalk_non_epi profile with the value

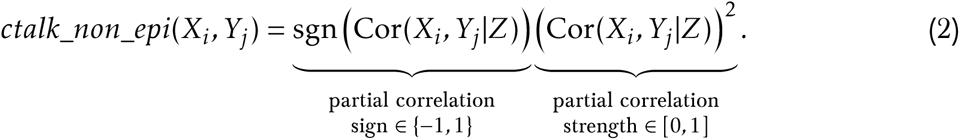

### Statistical significance assessment

The statistical significance of the partial correlation values, necessary for constructing the ctalk_non_epi profiles, was estimated using Fisher’s z-transformation [57]. Multiple comparisons correction of the *p*-values associated with each ctalk_non_epi profile was performed using the Benjamini-Yekutieli method [58].

### Hierarchical cluster analysis of ctalk_non_epi and mRNA abundance profiles

The goal of this step was to evaluate the significant ctalk_non_epi-profile clusters (if any) in the phenograms (i.e., phenotypic similarity dendrograms) obtained from hierarchical cluster analysis (HCA). For each species, HCA was performed on (i) the ctalk_non_epi profiles of each cell type/sample (Fig. 2A, 2B, and 2C) and (ii) the log-transformed FPKM profiles (i.e., mRNA abundance) of each cell type/sample (Fig. S2A, S2B, and S2C). The metric used was the “correlation distance” (i.e., 1 – Cor (*x, y*)) and the cluster-linkage method chosen was UPGMA (Unweighted Pair Group Method with Arithmetic Mean) [59]. Cluster statistical significance was assessed as au (approximately unbiased) and bp (bootstrap probability) significance scores by nonparametric bootstrap resampling using the Pvclust [60] add-on package for the R software [61]. The number of bootstrap replicates in each analysis was 10,000.

Source datafiles and methodological steps in full detail can be found in the *SI Materials and Methods.*

## Acknowledgments

I wish to thank Angelika H. Hofmann for editing this paper into an English I could only dream of writing. I am especially grateful to John Tyler Dodge, horn soloist at the *Orquesta Filarmonica de Santiago,* for reviewing the English of the very first complete draft of this paper and his valuable questions, which pushed me to the limit of my abilities in the purpose of making the theoretical description self-explanatory. For reviewing this paper and their valuable comments I am indebted to Álvaro Glavic, Óscar M. Lazo, Inti Pedroso, Iván Sellés, and José Monserrat Neto. My thanks also extend to Alejandro Maass and Kenneth M. Weiss for their interest in this work and their valuable questions and to my anonymous colleagues who reviewed my grant proposal on behalf of FONDECYT. This work was funded by the FONDECYT postdoctoral grant 3140328.

## Copyright

The copyright holder for this preprint is the author. It is made available under a Creative Commons Attribution 4.0 International License (bioRxiv doi: 10.1101/016840). To view a copy of this license, visit http://creativecommons.org/licenses/by/4.0/.

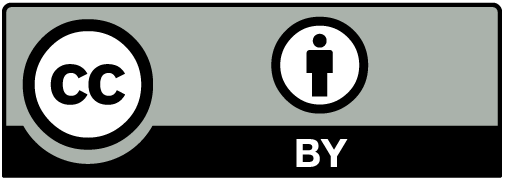

## Supplementary Information

**Fig. S1.**
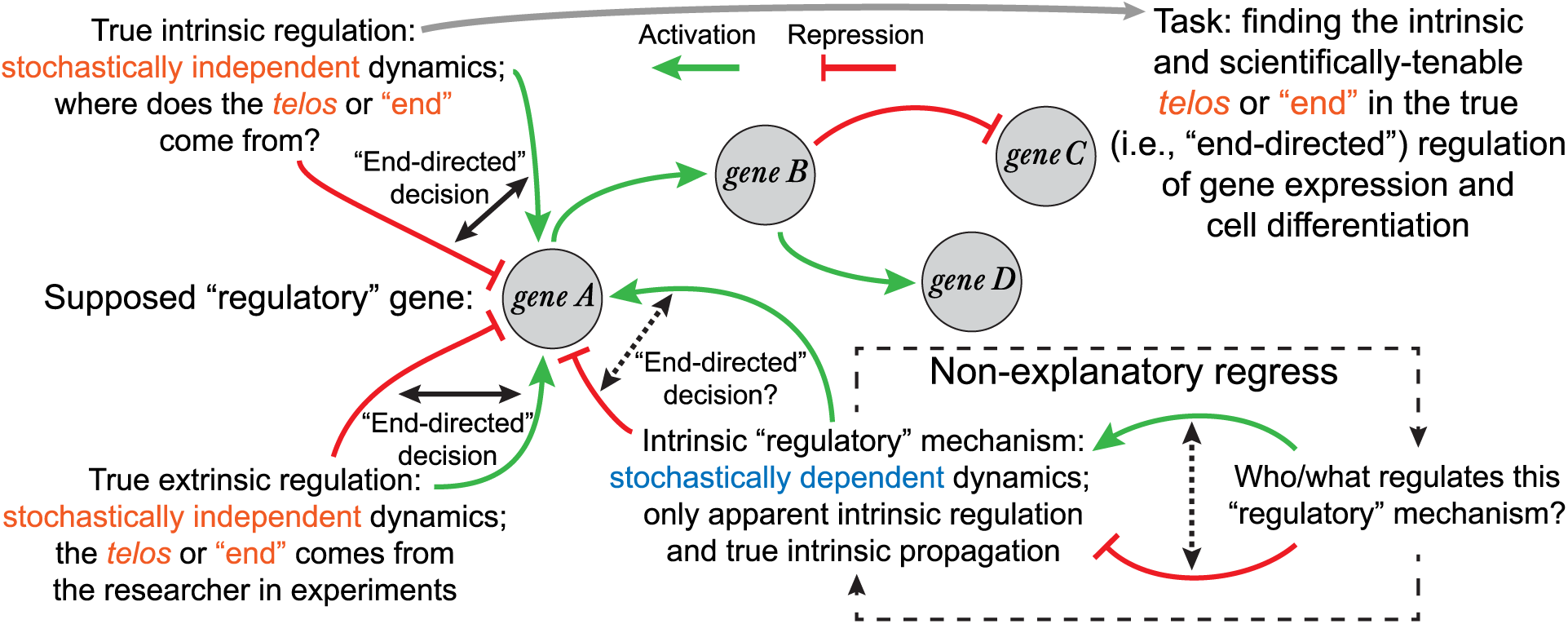
Some limitations of the epigenetic landscape model. This framework either falls into a non-explanatory regress when attempting to account for the intrinsic regulation of changes in gene expression for cell differentiation or uses “regulation” simply as a placeholder for what is only the propagation of such changes, bracketing true intrinsic regulation from further inquiry.

**Fig. S2.**
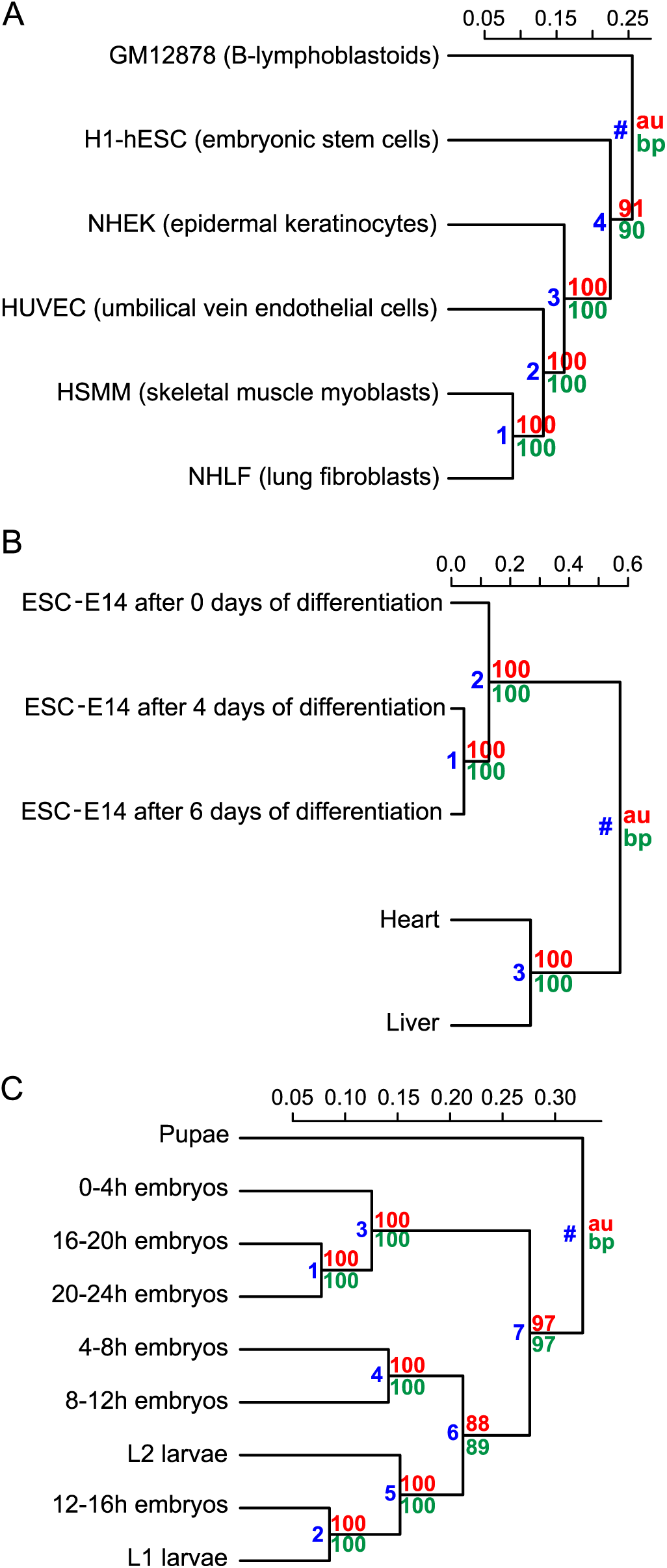
Hierarchical cluster analysis of mRNA abundance profiles. Organisms: *Homo sapiens* (A), *Mus musculus* (B), and *Drosophila melanogaster* (C). Metric: correlation (i.e., 1 – Cor (*x,y*)). Linkage method: UPGMA. Significance scores: au (approximately unbiased) and bp (bootstrap probability) [60]. Significant clusters were identified as those for which au and bp ≥ 95. Cluster identification numbers are in blue.

**Fig. S3.**
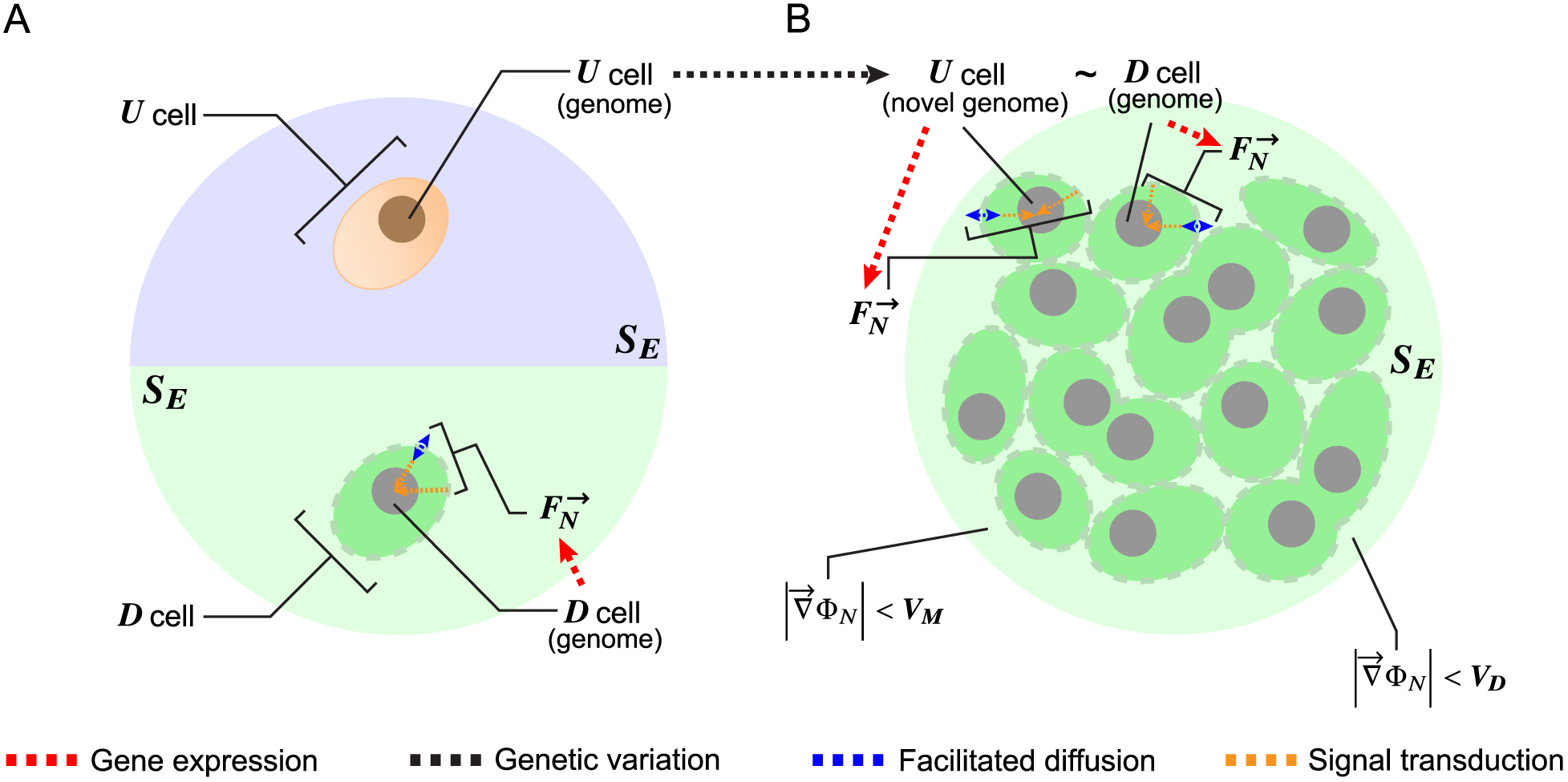
Necessary initial conditions for individuated multicellularity. (A, top) A cell of the unicellular and undifferentiated ancestor species *U*. (A, bottom) A primordial cell of the multicellular species *D*. (A, top to B, top) The necessary genetic change for individuated multicellularity occurs in the species *U*. (B, top) The similar and necessary alleles are now present in both species. (B, bottom) Cells proliferate but no significant 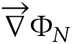 gradients form yet in *S_E_* and no differentiation is observed.

**Fig. S4.**
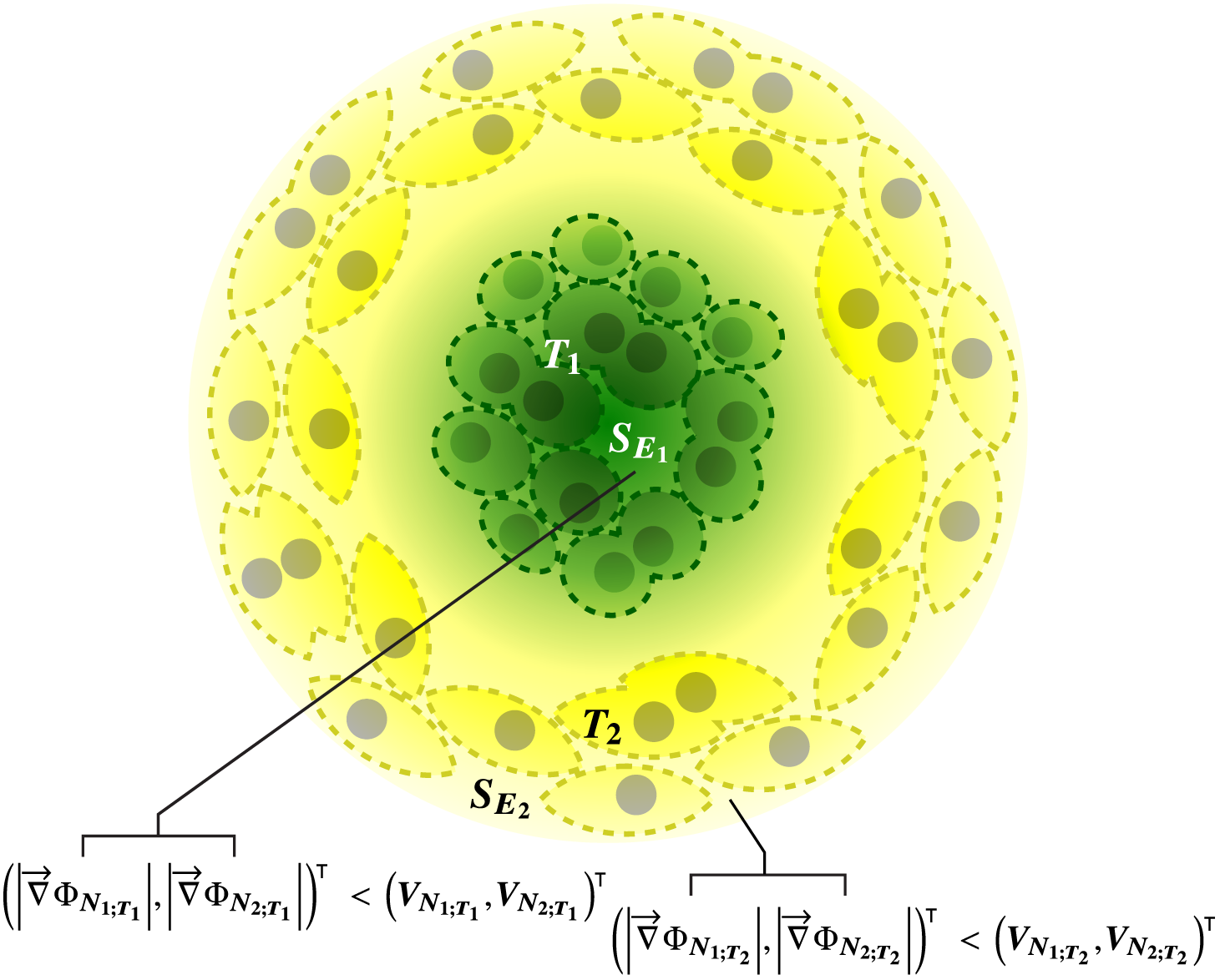
Novel cell types develop. Two distinct cell types T_1_ and T_2_ develop respectively in regions 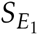 and 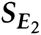 within *S_E_* characterized by a relative small 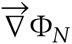 gradient magnitude, i.e., in extracellular regions of relative 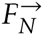 uniformity.

**Fig. S5.**
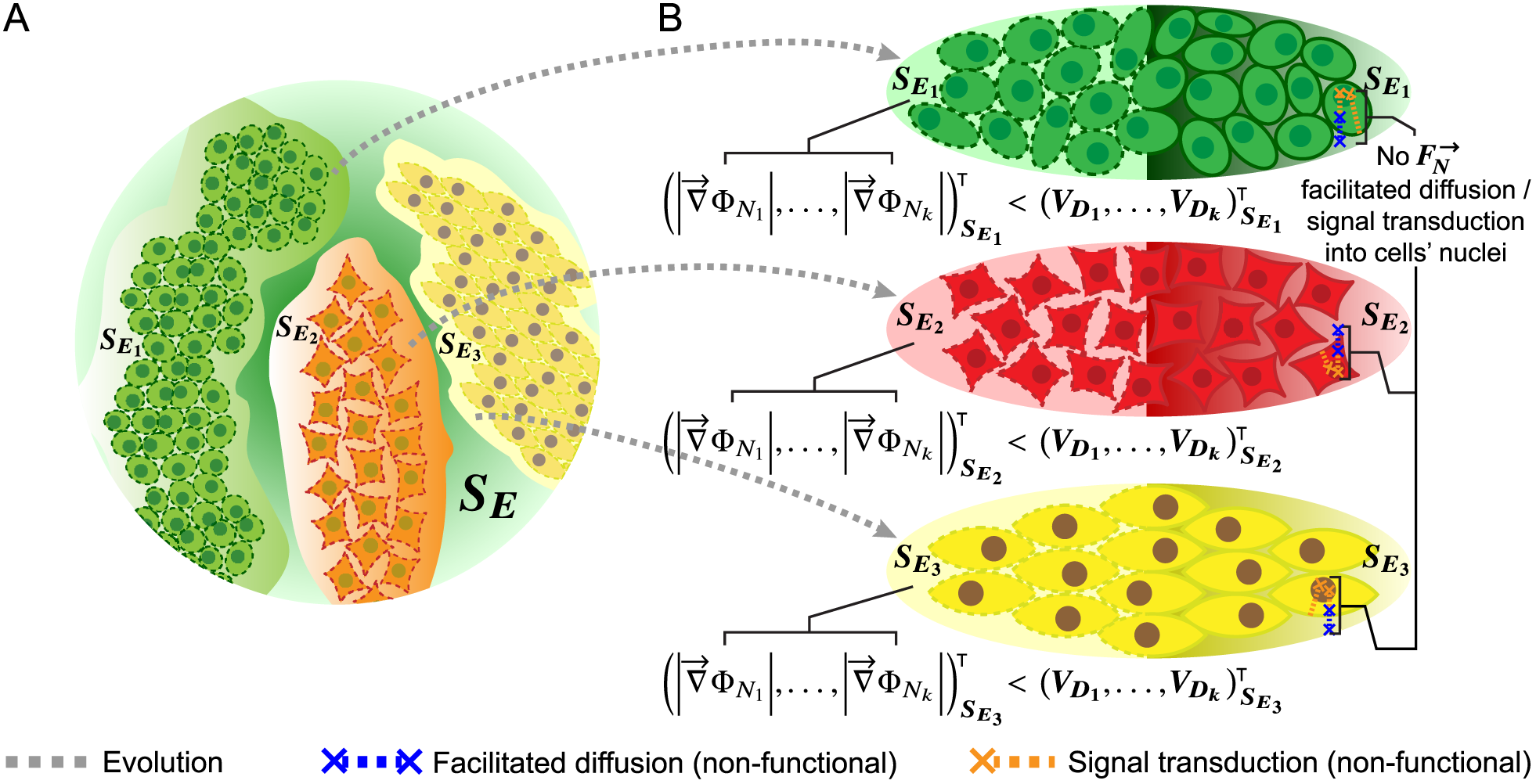
The evolution of cell types/tissues/organs and the “termination” of cell differentiation. (A) Cell types/tissues/organs evolve as emergent “blobs” of relatively small 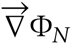 magnitude and then are shaped by evolution. (B) Cell differentiation stops when the 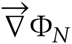 gradients dissipate (left), or when they cannot diffuse/be transduced into the cells’ nuclei (right).

**Table S1.** Estimation of a lower bound for the necessary cell-fate information capacity in the hermaphrodite *Caenorhabditis elegans* ontogeny.

| Count | N° |  |  |
| --- | --- | --- | --- |
| Cells generated | 1,090 |  |  |
| Deaths in the process | 131 |  |  |
| Final cells | 959 |  |  |
| Cell types developed | 19 |  |  |
| (Data source: WormAtlas website [18]) |  |  |  |
|  | Estimated as | N° (approx.) |  |
| Total divisions | $2^{\log_2(\text{cells\_generated}+1)} - 1$ | 2,179 | |
| Cell-fate divisions | $2^{\log_2(\text{cell\_types}+1)} - 1$ | 37 | |
| Non-cell-fate divisions | $\text{total\_divisions} - (\text{cell\_fate\_divisions} + \text{deaths})$ | 2,011 | |
| | Estimated as | $p$ | $-p \log_2 p$ |
| Cell death | $\text{deaths} / \text{total\_divisions}$ | 0.060 | 0.244 |
| Non-cell-fate division | $\text{non\_cell\_fate\_divisions} / \text{total\_divisions}$ | 0.923 | 0.107 |
| Cell-fate division | $\text{cell\_fate\_divisions} / \text{total\_divisions}$ | 0.017 | 0.1 |
| Uncertainty per division (Sum) |  |  | 0.451 |
|  | Estimated as | (bit) |  |
| Uncertainty to resolve (total) | $\text{uncertainty\_per\_division} \times \text{total\_divisions}$ | 983 | |

Note: germ line cells were excluded from the analysis.

### SI Corollaries

Corollaries, hypotheses and predictions (not involving falsifiability) that can be derived from the proposed theory include:

1. **Cell surface-to-volume ratio and the evolution and development of the extracellular matrix (ECM).** Under the predicted relationship between regenerative capacity and cells’ surface-to-volume ratio (see Part XII) neuron-shaped cells are expected to be the most difficult to regenerate. This is the developmental price to pay for an even higher-order, dynamically faster form of multicellular *self* that neurons make possible. On the other hand, glial cells (companions of neurons in the nervous tissue) have a smaller surface-to-volume ratio than neurons so they would support neurons by constraining to some extent the diffusion flux of Nanney’s extracellular propagators 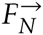 in the extracellular space *S_E_*. This hypothesis is supported by the fact that the cells serving as neural stem cells are ependymal cells [62], which are precisely those of the smallest surface-to-volume ratio in the neuroglia. It can thus be hypothesized that the extracellular matrix was not only shaped by evolution making it provide the cells biochemical and structural support but also developmental support, understood as fine-tuned differential constraints to the diffusion flux of 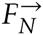 in *S_E_*. This necessary fine-tuning can underpin the observed relationship between stem cell fate and the topography and mechanical properties of the ECM (as reviewed in [63]).
2. **Natural developmental arrests or diapauses**. Natural diapauses (observable in arthropods [64] and some species of killifish [65]) are under this theory a metastable equilibrium state characterized by (i) the dissipation of the extracellular gradients 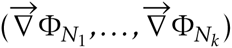 in *S_E_* under certain critical values because the otherwise relevant Nanney’s extracellular propagators 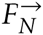 are no longer expressed or functional, or (ii) the inability of these gradients to modify Nanney’s constraints *C_N_* in the cells’ nuclei because the critical gene products for protein channels/carriers or membrane receptors/signal transducers are not expressed or non-functional. For example, if in some organism the function of the gene products critical for the facilitated diffusion/signal transduction of the current 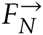 is temperature-dependent, then at that time development will enter a diapause given certain thermal conditions and resume when those conditions are lost.
3. 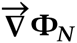 **gradients and tissue regeneration**. Under this theory it can be hypothesized that an important constraint driving the regenerative response to wounds is the magnitude of the gradient 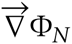 generated by the wound itself. This drive occurs because a wound creates an immediate, significant gradient at its edges. If different tissues of the same multicellular individual are compared, a significant negative correlation should be observable between the regenerative capacity after injury in a tissue and the average cell surface-to-volume ratio in that tissue, once controlling for average cell characteristic length. Evidence related to this corollary has been found already as extracellular H_2_O_2_ gradients mediating wound detection in zebrafish [66].
4. **Effects of microgravity on development**. In the last few decades a number of abnormal effects of microgravity on development-related phenomena have been described, including for mammal tissue culture [67], plant growth [68], human gene expression [69], cytoskeleton organization and general embryo development ([70] and references therein). I suggest that a key perturbation on development elicitable by microgravity is a significant alteration of the instantaneous 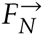 distribution in the extracellular space *S_E_*. This could be accounted for by changes in mass transfer dynamics as evidence for changes in the diffusion of miscible fluids suggest [71], and/or a significant density difference between the extracellular space *S_E_* and the cells (i.e., bulk flow).
5. **Why plant seeds need water**. It is a well-known fact that plant seeds only need certain initial water intake to be released from dormancy and begin to germinate with no additional nutrient supply until they are able to photosynthesize. Whereas this specific requirement of water has been associated to embryo expansion and metabolic activation of the seeds [72, 73], I submit that it is also associated to the fundamental need for a proper medium in *S_E_* where the critical gradients 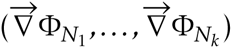 can form. These gradients would be in turn required for the intrinsic regulation of the asymmetric divisions already shown critical for cell differentiation in plants [74].
6. **The (lost) multicellular *telos* of cancer cells**. Cancer cells proliferate faster than normal cells regardless of the needs of their host, which is frequently killed once cancer metastasizes. It can thus be hypothesized under this theory that a *necessary* condition for the onset of cancer is that the intrinsic, higher-order constraint described here as the *telos* of individuated multicellularity has dissipated in cancer cells. In turn, such dissipation could only be accounted for by (i) the otherwise functional Nanney’s extracellular propagators 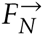 lacking their defining ability to elicit changes in Nanney’s constraints *C_N_* within other cells’ nuclei, (ii) no elements of the molecular nuclear phenotype being specifiable as Waddington’s embodiers *F_W_* and Nanney’s embodiers *F_N_* at the same time (such as histone H3 modifications in the data analyzed), thus preventing the coupling across *S_E_* of the respective *C_W_* and *C_N_* constraints, and/or (iii) Waddington’s embodiers *F_W_* not being able to constrain any longer—via gene expression and function—the facilitated diffusion/signal transduction of 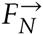 into the cells.

### SI Materials and Methods

#### Data collection

The genomic coordinates of all annotated RefSeq TSSs for the hg19 (*Homo sapiens*), mm9 (*Mus musculus*), and dm3 (*Drosophila melanogaster*) assemblies were downloaded from the UCSC (University of California, Santa Cruz) database [48]. Publicly available tandem datafiles of ChIP-seq (comprising 1×36 bp, 1×50 bp, and 1×75 bp reads, depending on the data series) on histone H3 modifications and RNA-seq (comprising 1×36 bp, 1×100 bp, and 2×75 bp reads, depending on the data series) for each analyzed cell sample in each species were downloaded from the ENCODE, modENCODE or the SRA (Sequence Read Archives) database of the National Center for Biotechnology Information [49, 50, 51, 52, 53, 54, 55].

The criteria for selecting cell type/cell sample datasets in each species were (i) excluding those associated to abnormal karyotypes and (ii) among the remaining datasets, choosing the group that maximizes the number of specific histone H3 modifications shared. Under these criteria, the cell type/sample datasets included in this work for computing *ctalk_non_epi* and mRNA abundance profiles were thus:

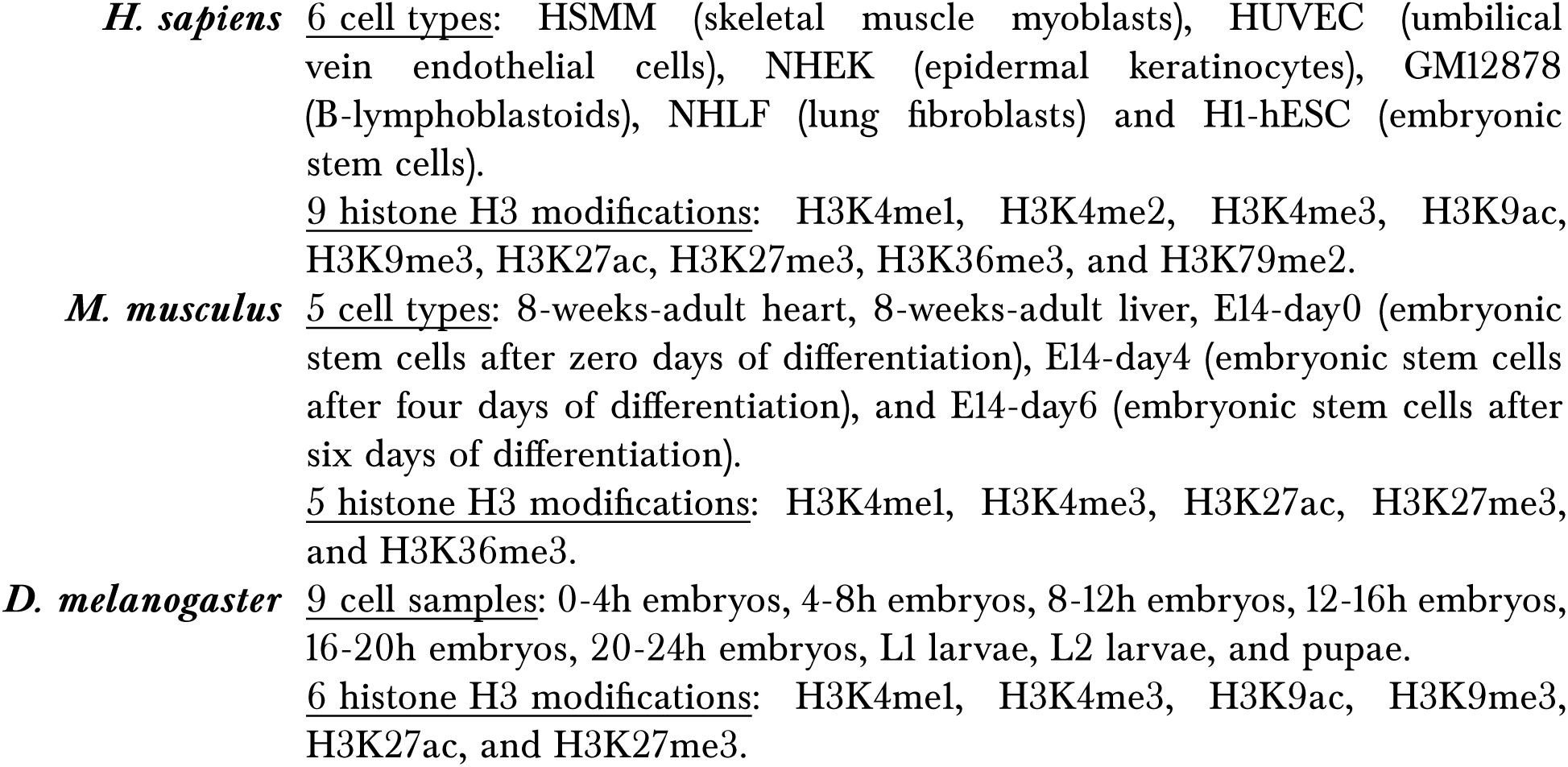

#### ChIP-seq read profiles and normalization

The first steps in the EFilter algorithm by Kumar *et al*.—which predicts mRNA levels in log-FPKM (fragments per transcript kilobase per million fragments mapped) with high accuracy (R~0.9) [25]—were used to generate ChIP-seq read signal profiles for the histone H3 modifications data. Namely, (i) dividing the genomic region from 2 kbp upstream to 4 kbp downstream of each TSS into 30 200-bp-long bins, in each of which ChIP-seq reads were later counted; (ii) dividing the read count signal for each bin by its corresponding control (Input/IgG) read density to minimize artifactual peaks; (iii) estimating this control read density within a 1-kbp window centered on each bin, if the 1-kbp window contained at least 20 reads. Otherwise, a 5-kbp window, or else a 10-kbp window was used if the control reads were less than 20. When the 10-kbp length was insufficient, a pseudo-count value of 20 reads per 10 kbp was set as the control read density. This implies that the denominator (i.e., control read density) is at least 0.4 reads per bin. When replicates were available, the measure of central tendency used was the median of the replicate read count values.

#### ChIP-seq read count processing

When the original format was SRA, each datafile was pre-processed with standard tools in the pipeline

~~~
fastq-dump → bwa aln [genome.fa]→ bwa samse → samtools view -bS -F 4 → samtools sort → samtools index
~~~

to generate its associated BAM (Binary Sequence Alignment/Map) and BAI (BAM Index) files. Otherwise, the tool

~~~
bedtools multicov -bams [file.bam] -bed [bins_and_controlwindows.bed]
~~~

was applied (excluding failed-QC reads and duplicate reads by default) directly on the original BAM file (the BAI file is required implicitly) to generate the corresponding read count file in BED (Browser Extensible Data) format.

#### RNA-seq data processing

The processed data were mRNA abundances in FPKM at RefSeq TSSs. When the original file format was GTF (Gene transfer Format) containing already FPKM values (as in the selected ENCODE RNA-seq datafiles for *H. sapiens*), those values were used directly in the analysis. When the original format was SAM (Sequence Alignment/Map), each datafile was pre-processed by first sorting it to generate then a BAM file using samtools view -bS. If otherwise the original format was BAM, mRNA levels at RefSeq TSSs were then calculated with FPKM as unit using *Cufflinks* [56] directly on the original file with the following three options:

~~~
-GTF-guide <reference_annotation.(gtf/gff)>
-frag-bias-correct <genome.fa>
-multi-read-correct
~~~

When the same TSS (i.e., same genomic coordinate and strand) displayed more than one identified transcript in the *Cufflinks* output, the respective FPKM values were added. When replicates were available the measure of central tendency used was the median of the replicate FPKM values.

For each of the three species, all TSS_def_—defined as those TSSs with (i) measured mRNA abundance (i.e., FPKM > 0) and (ii) measured histone modification level (i.e., at least one ChIP-seq read signal greater than zero among the 30 genomic bins) in all cell types/cell samples—were determined. The number of TSS_def_ found for each species were 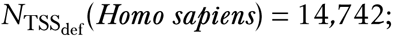 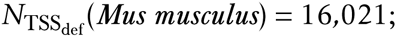 and 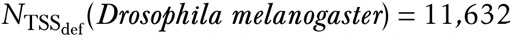. Then, for each cell type/cell sample, 30 genomic bins were defined and denoted by the distance (in bp) between their 5’-end and their respective TSS_def_ genomic coordinate: “−2000”, “−1800”, “−1600”, “−1400”, “−1200”, “−1000”, “−800”, “−600”, “−400”, “−200”, “0” (TSS_def_ or ‘+1’), “200”, “400”, “600”, “800”, “1000”, “1200”, “1400”, “1600”, “1800”, “2000”, “2200”, “2400”, “2600”, “2800”, “3000”, “3200”, “3400”, “3600”, and “3800”. Then, for each cell type/cell sample, a ChIP-seq read signal was computed for all bins in all TSS_def_ genomic regions (e.g., in the “−2000” bin of the *Homo sapiens* TSS with RefSeq ID: NM_001127328, H3K27ac_−2000 = 4.68 in H1-hESC stem cells). Data input tables, with *n_m_* being the number of histone H3 modifications comprised, were generated following this structure of rows and columns:

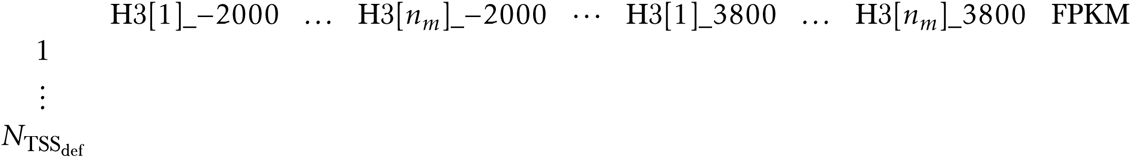

The tables were then written to the following data files:

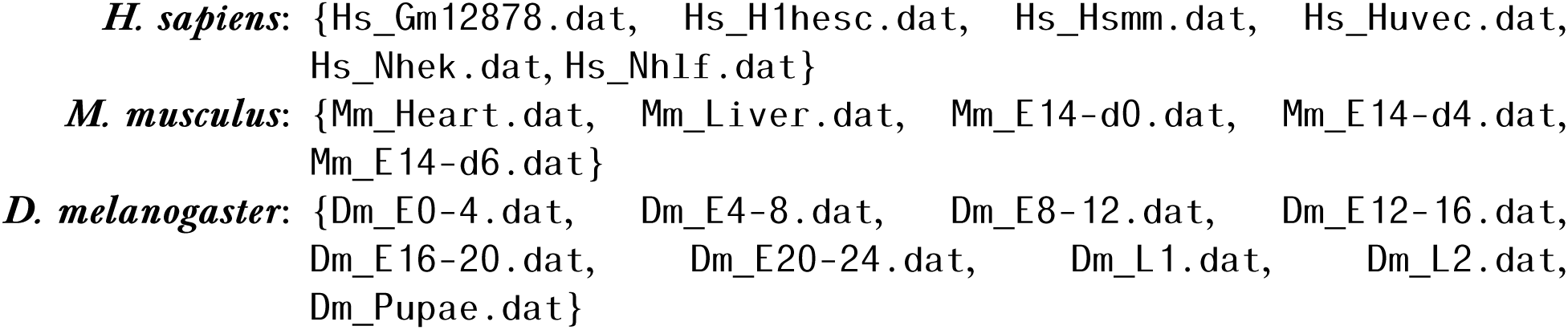

#### Computation of *ctalk_non_epi* profiles

If the variables *X_i_* (representing the signal for histone H3 modification *X* in the genomic bin *i ∈* {“− 2000”,...,”3800”}), *Y_j_* (representing the signal for histone H3 modification *Y* in the genomic bin *j ∈* {“− 2000”,...,”3800”}) and *Z* (representing log2-transformed FPKM values) are random variables, then the covariance of *X_i_* and *Y_j_* can be decomposed in terms of their linear relationship with *Z* as follows:

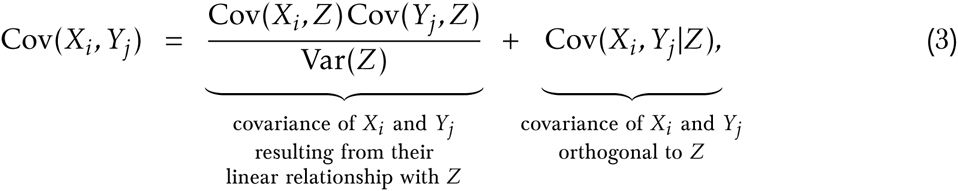

where the second summand *Cov(X_j_,Y_j_* |Z) is the partial covariance between *X_i_* and *Y_j_* given *Z*. It is easy to see that *Cov(X_i_,Y_j_* |Z) is a local approximation of Nanney’s constraints *C_N_* on histone H3 modifications, as anticipated in the preliminary theoretical definitions (a straightforward corollary is that Waddington’s constraints *C_W_* can in turn be approximated by 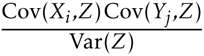). To make the *ctalk_non_epi* profiles comparable, however, Cov*(X_i_,Y_j_* |Z) values had to be normalized by the standard deviations of the residuals of *X_i_* and *Y_j_* with respect to *Z*. In other words, the partial correlation Cor(*X_i_,Y_j_* |Z) values were needed. Nevertheless, a correlation value does not have a straightforward interpretation, whereas its square—typically known as *coefficient of determination, strength of the correlation,* or simply *r*^2^—does: it represents the relative (i.e., fraction of) variance of one random variable explained by the other. For this reason, Cor(*X_i_*, *Y_j_*|Z)^2^ was used to represent the strength of the association, and then multiplied by the sign of the correlation to represent the direction of the association. Thus, after log_2_-transforming the *X_i_*, *Y_j_* and *Z* data, each pairwise combination of bin-specific histone H3 modifications *{X_i_, Y_j_*} contributed with the value

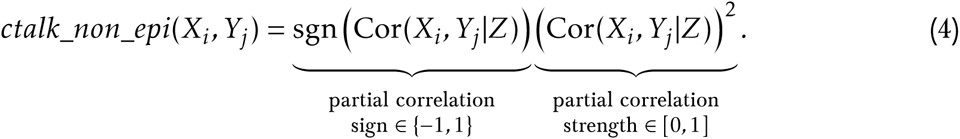

This implies that for each pairwise combination of histone H3 modifications {*X,Y*}, there are 30 (bins for *X*) × 30 (bins for *Y*) = 900 (bin-combination-specific *ctalk_non_epl* values). To increase the robustness of the analysis against the departures of the actual nucleosome distributions from the 30 × 200-bp bins model, the values were then sorted in descending order and placed in a 900-tuple.

For a cell type/cell sample from a species with data for *n_m_* histone H3 modifications, e.g., *n_m_(Mus musculus)* = 5, the length of the final *ctalk_non_epl* profile comprising all possible {*X, Y*} combinations would be 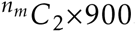. However, a final data filtering was performed.

The justification for this additional filtering was that some pairwise partial correlation values were expected to be strong and significant, which was later confirmed. Namely, (i) those involving the same histone H3 modification in the same amino acid residue (e.g., Cor(H3K9ac_–200,H3K9ac_–400|FPKM) > 0; Cor(H3K4me3_–200, H3K4me3_–200|FPKM) = 1), (ii) those involving a different type of histone H3 modification in the same amino acid residue (e.g., Cor(H3K27ac_-800,H3K27me3_–600|FPKM) < 0), and (iii) those involving the same type of histone H3 modification in the same amino acid residue (e.g., Cor(H3K4me2_–400,H3K4me3_–400|FPKM) > 0) in part because ChIP-antibody cross reactivity has been shown able to introduce artifacts on the accurate assessment of some histone-crosstalk associations [75, 76]. For these reasons, in each species all pairwise combinations of post-translational modifications involving the same amino acid residue in the H3 histone were then identified as “trivial” and excluded from the *ctalk_non_epl* profiles construction. e.g., for *Mus musculus* cell-type datasets the histone modifications comprised were H3K4me1, H3K4me3, H3K27ac, H3K27me3, and H3K36me3 (i.e., *n_m_* = 5), then the combinations H3K4me1-H3K4me3 and H3K27ac-H3K27me3 were filtered out. Therefore, the length of the *ctalk_non_epl* profiles for *Mus musculus* was (^5^C_2_ − 2) × 900 = 7,200.

#### Statistical significance assessment

The statistical significance of the partial correlation Cor(*X_i_,Y_j_ |Z*) values, necessary for constructing the *ctalk_non_epi* profiles, was estimated using Fisher’s z-transformation [57]. Under the null hypothesis Cor(*X_i_,Y_j_* |Z) = 0 the statistic 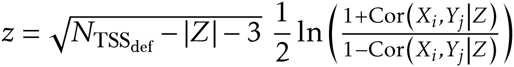 where 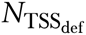 is the sample size and |*Z*| = 1 (i.e., one control variable), follows asymptotically a *N*(0,1) distribution. The *p*-values can then be computed easily using the *N*(0,1) probability function.

Multiple comparisons correction of the *p*-values associated with each *ctalk_non_epi* profile was performed using the Benjamini-Yekutieli method [58]. The parameter used was the number of all possible comparisons (i.e., *before* excluding “trivial” pairwise combinations of histone H3 modifications, to further increase the conservativeness of the correction): 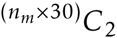. From the resulting q-values associated with each *ctalk_non_epi* profile an empirical cumulative distribution was obtained, which in turn was used to compute a threshold *t*. The value of *t* was optimized to be the maximum value such that within the *q*-values smaller than *t* it is expected less than 1 false-positive partial correlation value. Consequently, if *q*-value[*i*] ≥ *t* then the associated partial correlation value was identified as not significant (i.e., equal to zero) in the respective *ctalk_non_epi* profile.

#### Hierarchical cluster analysis of *ctalk_non_epi* and mRNA abundance profiles

The goal of this step was to evaluate the significant *ctalk_non_epi-profile* clusters (if any) in the phenograms (i.e., phenotypic similarity dendrograms) obtained from hierarchical cluster analysis (HCA). For each species, HCA was performed on (i) the *ctalk_non_epi* profiles of each cell type/sample (Figure 2A, 2B, and 2C) and (ii) the log2-transformed FPKM profiles (i.e., mRNA abundance) of each cell type/sample (Figure S2A, S2B, and S2C). Important to the HCA technique is the choice of a metric (for determining the distance between any two profiles) and a cluster-linkage method (for determining the distance between any two clusters).

Different ChIP-seq antibodies display differential binding affinities (with respect to different epitopes or even the same epitope, depending on the manufacturer) that are intrinsic and irrespective to the biological phenomenon of interest. For this reason, comparing directly the strengths (i.e., magnitudes) in the *ctalk_non_epi* profiles (e.g., using Euclidean distance as metric) is to introduce significant biases in the analysis. In contrast, the “correlation distance” metric—customarily used for comparing gene expression profiles—defined between any two profiles *pro*[*i*]*,pro*[*j*] as

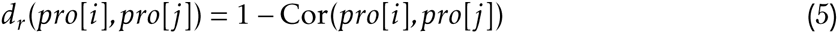

compares instead the “shape” of the profiles, hence it was the metric used here (as a consequence of what was highlighted previously, the “correlation distance” metric is also invariant under linear transformations of the profiles). On the other hand, the cluster-linkage method chosen was the UPGMA (Unweighted Pair Group Method with Arithmetic Mean) or “average” method in which the distance *D*(*A,B*) between any clusters *A* and *B* is defined as

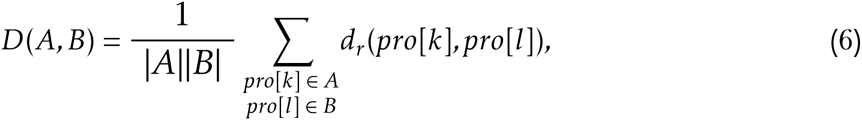

that is, the mean of all distances *d_r_*(*pro*[*k*]*,pro*[*l*]) such that *pro*[*k*] ∈ A and *pro*[*l*] ∈ *B* (this method was chosen because it has been shown to yield the highest cophenetic correlation values when using the “correlation distance” metric [59]). Cluster statistical significance was assessed as *au* (approximately unbiased) and *bp* (bootstrap probability) significance scores by nonparametric bootstrap resampling using the *Pvclust* [60] add-on package for the *R* software [61]. The number of bootstrap replicates in each analysis was 10,000.

#### Suitability of FPKM as unit of mRNA abundance

Previous research has pinpointed that FPKM may not always be an adequate unit of transcript abundance in differential expression studies. It was shown that, if transcript size distribution varies significantly among the samples, FPKM and RPKM (reads per kilobase of transcript per million reads mapped) may introduce significant biases. For this reason another abundance unit TPM (transcripts per million)—which is a linear transformation of the FPKM value for each sample—was proposed to overcome the limitation [77]. However, this issue was not a problem for the study because partial correlation, used to construct the *ctalk_non_epl* profiles, is invariant under linear transformations of the control variable *Z* (i.e., Cor(*X*, *Y\Z*) = Cor(*X,Y\aZ* + *b*) for any two scalars {*a,b*}). Importantly, this property also implies that *ctalk_non_epl* profiles are controlling not only for mRNA abundance but also for any other biological variable displaying a strong linear relationship with mRNA abundance (e.g., chromatin accessibility represented by DNase I hypersensitivity, as shown in [75]). Similarly, hierarchical clustering of mRNA abundance profiles is invariant under linear transformations of the profiles, because *Cor*(*Z_i_,Z_j_*) = Cor(*aZ_i_+b,cZ_j_+d*) (provided *ac >* 0).

#### SI Problems with current views

Since Ernst Haeckel’s “gastraea theory” [36], the most plausible models aimed to explain the evolution of individuated multicellularity are fundamentally divorced from the epigenetic model assumed to explain the self-regulatory dynamics underpinning individuated multicellularity. This is because Haeckel’s account and the models built upon it rely on the gradual specialization of same-species (or even different-species [78]) cell colonies or aggregations [36, 37, 38, 32, 39, 40, 41, 42] while the developmental process starts from one or a few primordial cells (zygotes, spores, or buds) or, in other words, “from the inside out”. Because individuated multicellularity is a single phenomenon whose evolution and self-regulation have been tackled by research under such divergent approaches, the resulting explanatory account is thus insufficiently substantiated as a whole.

Other, “non-epigenetic” hypotheses have been advanced aiming to explain the dynamics and/or informational requirements of cell-differentiation (which in turn could provide some hints on the evolution of multicellularity). One of them holds that spontaneous intercellular reaction-diffusion patterns are responsible for morphogenesis, and for cell differentiation as a consequence [29]. Although this model has been tested in terms of chemical differentiation of synthetic “cells” [79], it does not explain the critical relationship in which real differentiating/differentiated cells *serve* the individuated multicellular organism as a whole. Another hypothesis suggests that gene expression instability and stochasticity, in the context of external metabolic substrate gradients, create an intrinsic natural-selection-like mechanism able to drive the differentiation process [80]. A third “non-epigenetic” hypothesis is that cell fate decisions are the result of the characteristic coupling of gene expression and metabolism [81].

All of these accounts, however, fail to (i) explain how traits or dynamics that supposedly account for the transition to multicellularity or to cell differentiation have fundamentally analogous counterparts in nonindividuated multicellular or unicellular eukaryotic lineages, and/or (ii) account for the information required by developmental decisions for information and in the transition between strictly single-cell-related content to additional multicellular-individual-related content, and/or (iii) explain the reproducible and robust self-regulatory dynamics of gene expression for cell differentiation. These approaches also do not describe in an objective and unambiguous way the transition or difference between a highly complex or symbiotic cell population/aggregation and a differentiated multicellular individual, and they lack parsimony when encompassing both the evolution and self-regulation of individuated multicellularity. Neither are they falsifiable.

In contrast to these current hypotheses, the falsifiable theory proposed here regards the multicellular organism as a higher-order system that emerges from proliferating undifferentiated cells and *then* is subject to natural selection. The theoretical development in this work is not based on the substrate-based concept of irreducible emergence (fundamentally refuted by Jaegwon Kim [82, 83]) but instead converged from the strict *stochastically-independent-dynamics* condition argued in the introduction into what can be described as the constraint-based concept of emergence of unprecedented, higher-order teleological systems, pioneered in a broader perspective by Terrence Deacon in 2011 [33]. Importantly, this formulation of emergence does not build upon traditional concepts of *telos* or “final cause” but instead redefines the *telos* as a thermodynamically spontaneous, intrinsic constraint whose causal power is exerted at the present instant.

#### *Homo sapiens* source data of ChIP-seq on histone H3 modifications (BAM/BAI files) [50]

For downloading, the URL must be constructed by adding the following prefix to each file listed:

~~~
ftp://hgdownload.cse.ucsc.edu/goldenPath/hg19/encodeDCC/wgEncodeBroadHistone/
~~~

| Cell type | Antibody | GEO Accession | File URL suffix |
| --- | --- | --- | --- |
| GM12878 | H3K27ac | GSM733771 | wgEncodeBroadHistoneGm12878H3k27acStdA1nRep1.bam.bai |
| GM12878 | H3K27ac | GSM733771 | wgEncodeBroadHistoneGm12878H3k27acStdA1nRep1.bam |
| GM12878 | H3K27ac | GSM733771 | wgEncodeBroadHistoneGm12878H3k27acStdA1nRep2.bam.bai |
| GM12878 | H3K27ac | GSM733771 | wgEncodeBroadHistoneGm12878H3k27acStdA1nRep2.bam |
| GM12878 | H3K27me3 | GSM733758 | wgEncodeBroadHistoneGm12878H3k27me3StdA1nRep1.bam.bai |
| GM12878 | H3K27me3 | GSM733758 | wgEncodeBroadHistoneGm12878H3k27me3StdA1nRep1.bam |
| GM12878 | H3K27me3 | GSM733758 | wgEncodeBroadHistoneGm12878H3k27me3StdA1nRep2.bam.bai |
| GM12878 | H3K27me3 | GSM733758 | wgEncodeBroadHistoneGm12878H3k27me3StdA1nRep2.bam |
| GM12878 | H3K27me3 | GSM733758 | wgEncodeBroadHistoneGm12878H3k27me3StdA1nRep3V2.bam.bai |
| GM12878 | H3K27me3 | GSM733758 | wgEncodeBroadHistoneGm12878H3k27me3StdA1nRep3V2.bam |
| GM12878 | H3K36me3 | GSM733679 | wgEncodeBroadHistoneGm12878H3k36me3StdA1nRep1.bam.bai |
| GM12878 | H3K36me3 | GSM733679 | wgEncodeBroadHistoneGm12878H3k36me3StdA1nRep1.bam |
| GM12878 | H3K36me3 | GSM733679 | wgEncodeBroadHistoneGm12878H3k36me3StdA1nRep2.bam.bai |
| GM12878 | H3K36me3 | GSM733679 | wgEncodeBroadHistoneGm12878H3k36me3StdA1nRep2.bam |
| GM12878 | H3K4me1 | GSM733772 | wgEncodeBroadHistoneGm12878H3k4me1StdA1nRep2.bam.bai |
| GM12878 | H3K4me1 | GSM733772 | wgEncodeBroadHistoneGm12878H3k4me1StdA1nRep2.bam |
| GM12878 | H3K4me1 | GSM733772 | wgEncodeBroadHistoneGm12878H3k04me1StdA1nRep1V2.bam.bai |
| GM12878 | H3K4me1 | GSM733772 | wgEncodeBroadHistoneGm12878H3k04me1StdA1nRep1V2.bam |
| GM12878 | H3K4me2 | GSM733769 | wgEncodeBroadHistoneGm12878H3k4me2StdA1nRep1.bam.bai |
| GM12878 | H3K4me2 | GSM733769 | wgEncodeBroadHistoneGm12878H3k4me2StdA1nRep1.bam |
| GM12878 | H3K4me2 | GSM733769 | wgEncodeBroadHistoneGm12878H3k4me2StdA1nRep2.bam.bai |
| GM12878 | H3K4me2 | GSM733769 | wgEncodeBroadHistoneGm12878H3k4me2StdA1nRep2.bam |
| GM12878 | H3K4me3 | GSM733708 | wgEncodeBroadHistoneGm12878H3k04me3StdA1nRep2V2.bam.bai |
| GM12878 | H3K4me3 | GSM733708 | wgEncodeBroadHistoneGm12878H3k04me3StdA1nRep2V2.bam |
| GM12878 | H3K4me3 | GSM733708 | wgEncodeBroadHistoneGm12878H3k4me3StdA1nRep1.bam.bai |
| GM12878 | H3K4me3 | GSM733708 | wgEncodeBroadHistoneGm12878H3k4me3StdA1nRep1.bam |
| GM12878 | H3K79me2 | GSM733736 | wgEncodeBroadHistoneGm12878H3k79me2StdA1nRep1.bam.bai |
| GM12878 | H3K79me2 | GSM733736 | wgEncodeBroadHistoneGm12878H3k79me2StdA1nRep1.bam |
| GM12878 | H3K79me2 | GSM733736 | wgEncodeBroadHistoneGm12878H3k79me2StdA1nRep2.bam.bai |
| GM12878 | H3K79me2 | GSM733736 | wgEncodeBroadHistoneGm12878H3k79me2StdA1nRep2.bam |
| GM12878 | H3K9ac | GSM733677 | wgEncodeBroadHistoneGm12878H3k9acStdA1nRep1.bam.bai |
| GM12878 | H3K9ac | GSM733677 | wgEncodeBroadHistoneGm12878H3k9acStdA1nRep1.bam |
| GM12878 | H3K9ac | GSM733677 | wgEncodeBroadHistoneGm12878H3k9acStdA1nRep2.bam.bai |
| GM12878 | H3K9ac | GSM733677 | wgEncodeBroadHistoneGm12878H3k9acStdA1nRep2.bam |
| GM12878 | H3K9me3 | GSM733664 | wgEncodeBroadHistoneGm12878H3k9me3StdA1nRep1.bam.bai |
| GM12878 | H3K9me3 | GSM733664 | wgEncodeBroadHistoneGm12878H3k9me3StdA1nRep1.bam |
| GM12878 | H3K9me3 | GSM733664 | wgEncodeBroadHistoneGm12878H3k9me3StdA1nRep2.bam.bai |
| GM12878 | H3K9me3 | GSM733664 | wgEncodeBroadHistoneGm12878H3k9me3StdA1nRep2.bam |
| GM12878 | H3K9me3 | GSM733664 | wgEncodeBroadHistoneGm12878H3k9me3StdA1nRep3.bam.bai |
| GM12878 | H3K9me3 | GSM733664 | wgEncodeBroadHistoneGm12878H3k9me3StdA1nRep3.bam |
| GM12878 | Input | GSM733742 | wgEncodeBroadHistoneGm12878ControlStdA1nRep1.bam.bai |
| GM12878 | Input | GSM733742 | wgEncodeBroadHistoneGm12878ControlStdA1nRep1.bam |
| GM12878 | Input | GSM733742 | wgEncodeBroadHistoneGm12878ControlStdA1nRep2.bam.bai |
| GM12878 | Input | GSM733742 | wgEncodeBroadHistoneGm12878ControlStdA1nRep2.bam |
| Hi-hESC | H3K27ac | GSM733718 | wgEncodeBroadHistoneH1hescH3k27acStdA1nRep1.bam.bai |

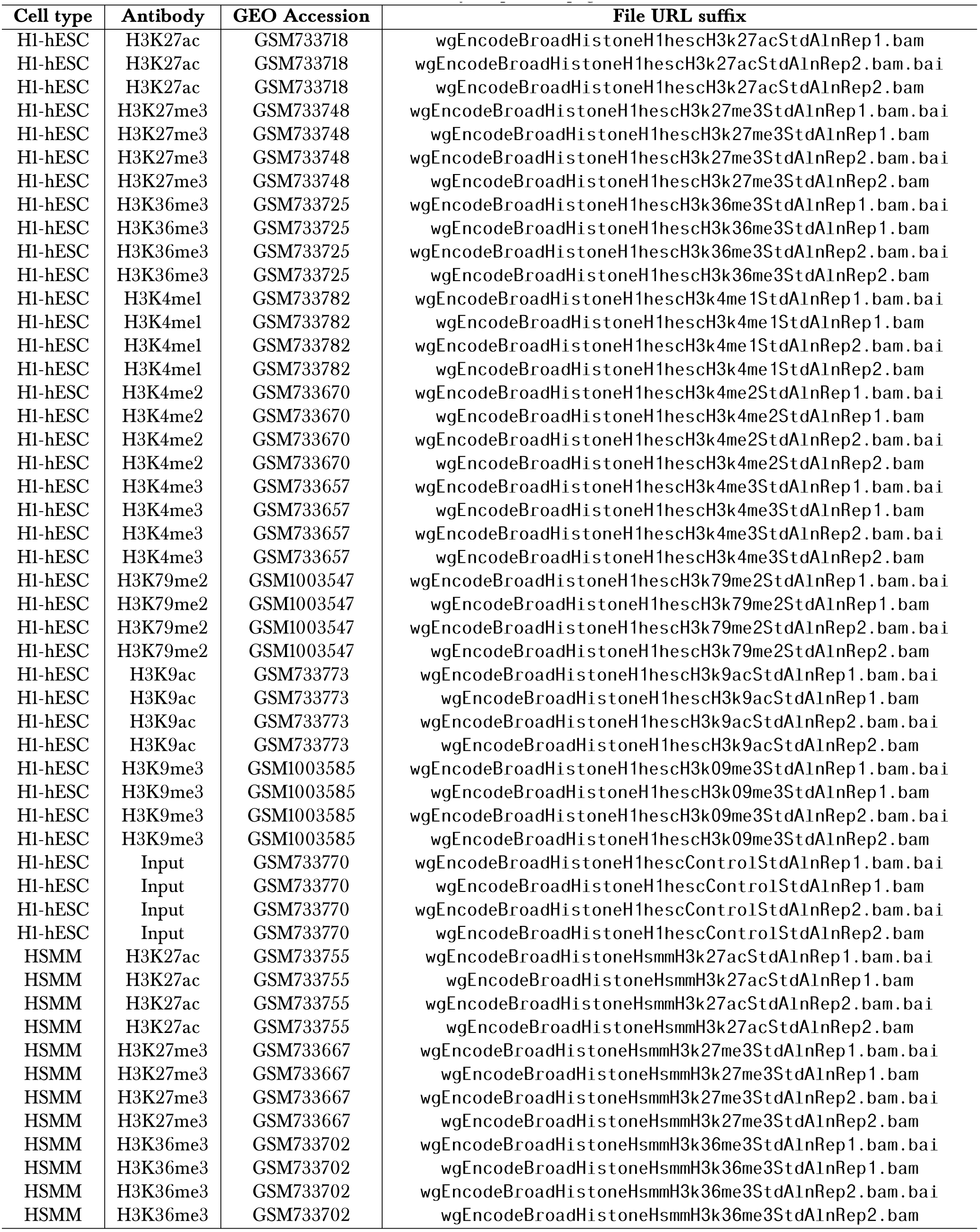

| Cell type | Antibody | GEO Accession | File URL suffix |
| --- | --- | --- | --- |
| Hi-hESC | H3K27ac | GSM733718 | wgEncodeBroadHistoneH1hesch3k27acStdA1nRep1.bam |
| Hi-hESC | H3K27ac | GSM733718 | wgEncodeBroadHistoneH1hesch3k27acStdA1nRep2.bam.bai |
| Hi-hESC | H3K27ac | GSM733718 | wgEncodeBroadHistoneH1hesch3k27acStdA1nRep2.bam |
| Hi-hESC | H3K27me3 | GSM733748 | wgEncodeBroadHistoneH1hesch3k27me3StdA1nRep1.bam.bai |
| Hi-hESC | H3K27me3 | GSM733748 | wgEncodeBroadHistoneH1hesch3k27me3StdA1nRep1.bam |
| Hi-hESC | H3K27me3 | GSM733748 | wgEncodeBroadHistoneH1hesch3k27me3StdA1nRep2.bam.bai |
| Hi-hESC | H3K27me3 | GSM733748 | wgEncodeBroadHistoneH1hesch3k27me3StdA1nRep2.bam |
| Hi-hESC | H3K36me3 | GSM733725 | wgEncodeBroadHistoneH1hesch3k36me3StdA1nRep1.bam.bai |
| Hi-hESC | H3K36me3 | GSM733725 | wgEncodeBroadHistoneH1hesch3k36me3StdA1nRep1.bam |
| Hi-hESC | H3K36me3 | GSM733725 | wgEncodeBroadHistoneH1hesch3k36me3StdA1nRep2.bam.bai |
| Hi-hESC | H3K36me3 | GSM733725 | wgEncodeBroadHistoneH1hesch3k36me3StdA1nRep2.bam |
| Hi-hESC | H3K4me1 | GSM733782 | wgEncodeBroadHistoneH1hesch3k4me1StdA1nRep1.bam.bai |
| Hi-hESC | H3K4me1 | GSM733782 | wgEncodeBroadHistoneH1hesch3k4me1StdA1nRep1.bam |
| Hi-hESC | H3K4me1 | GSM733782 | wgEncodeBroadHistoneH1hesch3k4me1StdA1nRep2.bam.bai |
| Hi-hESC | H3K4me1 | GSM733782 | wgEncodeBroadHistoneH1hesch3k4me1StdA1nRep2.bam |
| Hi-hESC | H3K4me2 | GSM733670 | wgEncodeBroadHistoneH1hesch3k4me2StdA1nRep1.bam.bai |
| Hi-hESC | H3K4me2 | GSM733670 | wgEncodeBroadHistoneH1hesch3k4me2StdA1nRep1.bam |
| Hi-hESC | H3K4me2 | GSM733670 | wgEncodeBroadHistoneH1hesch3k4me2StdA1nRep2.bam.bai |
| Hi-hESC | H3K4me2 | GSM733670 | wgEncodeBroadHistoneH1hesch3k4me2StdA1nRep2.bam |
| Hi-hESC | H3K4me3 | GSM733657 | wgEncodeBroadHistoneH1hesch3k4me3StdA1nRep1.bam.bai |
| Hi-hESC | H3K4me3 | GSM733657 | wgEncodeBroadHistoneH1hesch3k4me3StdA1nRep1.bam |
| Hi-hESC | H3K4me3 | GSM733657 | wgEncodeBroadHistoneH1hesch3k4me3StdA1nRep2.bam.bai |
| Hi-hESC | H3K4me3 | GSM733657 | wgEncodeBroadHistoneH1hesch3k4me3StdA1nRep2.bam |
| Hi-hESC | H3K79me2 | GSM1003547 | wgEncodeBroadHistoneH1hesch3k79me2StdA1nRep1.bam.bai |
| Hi-hESC | H3K79me2 | GSM1003547 | wgEncodeBroadHistoneH1hesch3k79me2StdA1nRep1.bam |
| Hi-hESC | H3K79me2 | GSM1003547 | wgEncodeBroadHistoneH1hesch3k79me2StdA1nRep2.bam.bai |
| Hi-hESC | H3K79me2 | GSM1003547 | wgEncodeBroadHistoneH1hesch3k79me2StdA1nRep2.bam |
| Hi-hESC | H3K9ac | GSM733773 | wgEncodeBroadHistoneH1hesch3k9acStdA1nRep1.bam.bai |
| Hi-hESC | H3K9ac | GSM733773 | wgEncodeBroadHistoneH1hesch3k9acStdA1nRep1.bam |
| Hi-hESC | H3K9ac | GSM733773 | wgEncodeBroadHistoneH1hesch3k9acStdA1nRep2.bam.bai |
| Hi-hESC | H3K9ac | GSM733773 | wgEncodeBroadHistoneH1hesch3k9acStdA1nRep2.bam |
| Hi-hESC | H3K9me3 | GSM1003585 | wgEncodeBroadHistoneH1hesch3k09me3StdA1nRep1.bam.bai |
| Hi-hESC | H3K9me3 | GSM1003585 | wgEncodeBroadHistoneH1hesch3k09me3StdA1nRep1.bam |
| Hi-hESC | H3K9me3 | GSM1003585 | wgEncodeBroadHistoneH1hesch3k09me3StdA1nRep2.bam.bai |
| Hi-hESC | H3K9me3 | GSM1003585 | wgEncodeBroadHistoneH1hesch3k09me3StdA1nRep2.bam |
| Hi-hESC | Input | GSM733770 | wgEncodeBroadHistoneH1hescontrolStdA1nRep1.bam.bai |
| Hi-hESC | Input | GSM733770 | wgEncodeBroadHistoneH1hescontrolStdA1nRep1.bam |
| Hi-hESC | Input | GSM733770 | wgEncodeBroadHistoneH1hescontrolStdA1nRep2.bam.bai |
| Hi-hESC | Input | GSM733770 | wgEncodeBroadHistoneH1hescontrolStdA1nRep2.bam |
| HSMM | H3K27ac | GSM733755 | wgEncodeBroadHistoneHsmmH3k27acStdA1nRep1.bam.bai |
| HSMM | H3K27ac | GSM733755 | wgEncodeBroadHistoneHsmmH3k27acStdA1nRep1.bam |
| HSMM | H3K27ac | GSM733755 | wgEncodeBroadHistoneHsmmH3k27acStdA1nRep2.bam.bai |
| HSMM | H3K27ac | GSM733755 | wgEncodeBroadHistoneHsmmH3k27acStdA1nRep2.bam |
| HSMM | H3K27me3 | GSM733667 | wgEncodeBroadHistoneHsmmH3k27me3StdA1nRep1.bam.bai |
| HSMM | H3K27me3 | GSM733667 | wgEncodeBroadHistoneHsmmH3k27me3StdA1nRep1.bam |
| HSMM | H3K27me3 | GSM733667 | wgEncodeBroadHistoneHsmmH3k27me3StdA1nRep2.bam.bai |
| HSMM | H3K27me3 | GSM733667 | wgEncodeBroadHistoneHsmmH3k27me3StdA1nRep2.bam |
| HSMM | H3K36me3 | GSM733702 | wgEncodeBroadHistoneHsmmH3k36me3StdA1nRep1.bam.bai |
| HSMM | H3K36me3 | GSM733702 | wgEncodeBroadHistoneHsmmH3k36me3StdA1nRep1.bam |
| HSMM | H3K36me3 | GSM733702 | wgEncodeBroadHistoneHsmmH3k36me3StdA1nRep2.bam.bai |
| HSMM | H3K36me3 | GSM733702 | wgEncodeBroadHistoneHsmmH3k36me3StdA1nRep2.bam |

| Cell type | Antibody | GEO Accession | File URL suffix |
| --- | --- | --- | --- |
| HSMM | H3K4me1 | GSM733761 | wgEncodeBroadHistoneHsmmH3k4me1StdA1nRep1.bam.bai |
| HSMM | H3K4me1 | GSM733761 | wgEncodeBroadHistoneHsmmH3k4me1StdA1nRep1.bam |
| HSMM | H3K4me1 | GSM733761 | wgEncodeBroadHistoneHsmmH3k4me1StdA1nRep2.bam.bai |
| HSMM | H3K4me1 | GSM733761 | wgEncodeBroadHistoneHsmmH3k4me1StdA1nRep2.bam |
| HSMM | H3K4me2 | GSM733768 | wgEncodeBroadHistoneHsmmH3k4me2StdA1nRep1.bam.bai |
| HSMM | H3K4me2 | GSM733768 | wgEncodeBroadHistoneHsmmH3k4me2StdA1nRep1.bam |
| HSMM | H3K4me2 | GSM733768 | wgEncodeBroadHistoneHsmmH3k4me2StdA1nRep2.bam.bai |
| HSMM | H3K4me2 | GSM733768 | wgEncodeBroadHistoneHsmmH3k4me2StdA1nRep2.bam |
| HSMM | H3K4me3 | GSM733637 | wgEncodeBroadHistoneHsmmH3k4me3StdA1nRep1.bam.bai |
| HSMM | H3K4me3 | GSM733637 | wgEncodeBroadHistoneHsmmH3k4me3StdA1nRep1.bam |
| HSMM | H3K4me3 | GSM733637 | wgEncodeBroadHistoneHsmmH3k4me3StdA1nRep2.bam.bai |
| HSMM | H3K4me3 | GSM733637 | wgEncodeBroadHistoneHsmmH3k4me3StdA1nRep2.bam |
| HSMM | H3K79me2 | GSM733741 | wgEncodeBroadHistoneHsmmH3k79me2StdA1nRep1.bam.bai |
| HSMM | H3K79me2 | GSM733741 | wgEncodeBroadHistoneHsmmH3k79me2StdA1nRep1.bam |
| HSMM | H3K79me2 | GSM733741 | wgEncodeBroadHistoneHsmmH3k79me2StdA1nRep2.bam.bai |
| HSMM | H3K79me2 | GSM733741 | wgEncodeBroadHistoneHsmmH3k79me2StdA1nRep2.bam |
| HSMM | H3K9ac | GSM733775 | wgEncodeBroadHistoneHsmmH3k9acStdA1nRep1.bam.bai |
| HSMM | H3K9ac | GSM733775 | wgEncodeBroadHistoneHsmmH3k9acStdA1nRep1.bam |
| HSMM | H3K9ac | GSM733775 | wgEncodeBroadHistoneHsmmH3k9acStdA1nRep2.bam.bai |
| HSMM | H3K9ac | GSM733775 | wgEncodeBroadHistoneHsmmH3k9acStdA1nRep2.bam |
| HSMM | H3K9me3 | GSM733730 | wgEncodeBroadHistoneHsmmH3k9me3StdA1nRep1.bam.bai |
| HSMM | H3K9me3 | GSM733730 | wgEncodeBroadHistoneHsmmH3k9me3StdA1nRep1.bam |
| HSMM | H3K9me3 | GSM733730 | wgEncodeBroadHistoneHsmmH3k9me3StdA1nRep2.bam.bai |
| HSMM | H3K9me3 | GSM733730 | wgEncodeBroadHistoneHsmmH3k9me3StdA1nRep2.bam |
| HSMM | Input | GSM733663 | wgEncodeBroadHistoneHsmmControlStdA1nRep1.bam.bai |
| HSMM | Input | GSM733663 | wgEncodeBroadHistoneHsmmControlStdA1nRep1.bam |
| HSMM | Input | GSM733663 | wgEncodeBroadHistoneHsmmControlStdA1nRep2.bam.bai |
| HSMM | Input | GSM733663 | wgEncodeBroadHistoneHsmmControlStdA1nRep2.bam |
| HUVEC | H3K27ac | GSM733691 | wgEncodeBroadHistoneHuvecH3k27acStdA1nRep1.bam.bai |
| HUVEC | H3K27ac | GSM733691 | wgEncodeBroadHistoneHuvecH3k27acStdA1nRep1.bam |
| HUVEC | H3K27ac | GSM733691 | wgEncodeBroadHistoneHuvecH3k27acStdA1nRep2.bam.bai |
| HUVEC | H3K27ac | GSM733691 | wgEncodeBroadHistoneHuvecH3k27acStdA1nRep2.bam |
| HUVEC | H3K27ac | GSM733691 | wgEncodeBroadHistoneHuvecH3k27acStdA1nRep3.bam.bai |
| HUVEC | H3K27ac | GSM733691 | wgEncodeBroadHistoneHuvecH3k27acStdA1nRep3.bam |
| HUVEC | H3K27me3 | GSM733688 | wgEncodeBroadHistoneHuvecH3k27me3StdA1nRep1.bam.bai |
| HUVEC | H3K27me3 | GSM733688 | wgEncodeBroadHistoneHuvecH3k27me3StdA1nRep1.bam |
| HUVEC | H3K27me3 | GSM733688 | wgEncodeBroadHistoneHuvecH3k27me3StdA1nRep2.bam.bai |
| HUVEC | H3K27me3 | GSM733688 | wgEncodeBroadHistoneHuvecH3k27me3StdA1nRep2.bam |
| HUVEC | H3K36me3 | GSM733757 | wgEncodeBroadHistoneHuvecH3k36me3StdA1nRep1.bam.bai |
| HUVEC | H3K36me3 | GSM733757 | wgEncodeBroadHistoneHuvecH3k36me3StdA1nRep1.bam |
| HUVEC | H3K36me3 | GSM733757 | wgEncodeBroadHistoneHuvecH3k36me3StdA1nRep2.bam.bai |
| HUVEC | H3K36me3 | GSM733757 | wgEncodeBroadHistoneHuvecH3k36me3StdA1nRep2.bam |
| HUVEC | H3K36me3 | GSM733757 | wgEncodeBroadHistoneHuvecH3k36me3StdA1nRep3.bam.bai |
| HUVEC | H3K36me3 | GSM733757 | wgEncodeBroadHistoneHuvecH3k36me3StdA1nRep3.bam |
| HUVEC | H3K4me1 | GSM733690 | wgEncodeBroadHistoneHuvecH3k4me1StdA1nRep1.bam.bai |
| HUVEC | H3K4me1 | GSM733690 | wgEncodeBroadHistoneHuvecH3k4me1StdA1nRep1.bam |
| HUVEC | H3K4me1 | GSM733690 | wgEncodeBroadHistoneHuvecH3k4me1StdA1nRep2.bam.bai |
| HUVEC | H3K4me1 | GSM733690 | wgEncodeBroadHistoneHuvecH3k4me1StdA1nRep2.bam |
| HUVEC | H3K4me1 | GSM733690 | wgEncodeBroadHistoneHuvecH3k4me1StdA1nRep3.bam.bai |
| HUVEC | H3K4me1 | GSM733690 | wgEncodeBroadHistoneHuvecH3k4me1StdA1nRep3.bam |
| HUVEC | H3K4me2 | GSM733683 | wgEncodeBroadHistoneHuvecH3k4me2StdA1nRep1.bam.bai |

| Cell type | Antibody | GEO Accession | File URL suffix |
| --- | --- | --- | --- |
| HUVEC | H3K4me2 | GSM733683 | wgEncodeBroadHistoneHuvecH3k4me2StdA1nRep1.bam |
| HUVEC | H3K4me2 | GSM733683 | wgEncodeBroadHistoneHuvecH3k4me2StdA1nRep2.bam.bai |
| HUVEC | H3K4me2 | GSM733683 | wgEncodeBroadHistoneHuvecH3k4me2StdA1nRep2.bam |
| HUVEC | H3K4me3 | GSM733673 | wgEncodeBroadHistoneHuvecH3k4me3StdA1nRep1.bam.bai |
| HUVEC | H3K4me3 | GSM733673 | wgEncodeBroadHistoneHuvecH3k4me3StdA1nRep1.bam |
| HUVEC | H3K4me3 | GSM733673 | wgEncodeBroadHistoneHuvecH3k4me3StdA1nRep2.bam.bai |
| HUVEC | H3K4me3 | GSM733673 | wgEncodeBroadHistoneHuvecH3k4me3StdA1nRep2.bam |
| HUVEC | H3K4me3 | GSM733673 | wgEncodeBroadHistoneHuvecH3k4me3StdA1nRep3.bam.bai |
| HUVEC | H3K4me3 | GSM733673 | wgEncodeBroadHistoneHuvecH3k4me3StdA1nRep3.bam |
| HUVEC | H3K79me2 | GSM1003555 | wgEncodeBroadHistoneHuvecH3k79me2A1nRep1.bam.bai |
| HUVEC | H3K79me2 | GSM1003555 | wgEncodeBroadHistoneHuvecH3k79me2A1nRep1.bam |
| HUVEC | H3K79me2 | GSM1003555 | wgEncodeBroadHistoneHuvecH3k79me2A1nRep2.bam.bai |
| HUVEC | H3K79me2 | GSM1003555 | wgEncodeBroadHistoneHuvecH3k79me2A1nRep2.bam |
| HUVEC | H3K9ac | GSM733735 | wgEncodeBroadHistoneHuvecH3k9acStdA1nRep1.bam.bai |
| HUVEC | H3K9ac | GSM733735 | wgEncodeBroadHistoneHuvecH3k9acStdA1nRep1.bam |
| HUVEC | H3K9ac | GSM733735 | wgEncodeBroadHistoneHuvecH3k9acStdA1nRep2.bam.bai |
| HUVEC | H3K9ac | GSM733735 | wgEncodeBroadHistoneHuvecH3k9acStdA1nRep2.bam |
| HUVEC | H3K9ac | GSM733735 | wgEncodeBroadHistoneHuvecH3k9acStdA1nRep3.bam.bai |
| HUVEC | H3K9ac | GSM733735 | wgEncodeBroadHistoneHuvecH3k9acStdA1nRep3.bam |
| HUVEC | H3K9me3 | GSM1003517 | wgEncodeBroadHistoneHuvecH3k09me3A1nRep1.bam.bai |
| HUVEC | H3K9me3 | GSM1003517 | wgEncodeBroadHistoneHuvecH3k09me3A1nRep1.bam |
| HUVEC | H3K9me3 | GSM1003517 | wgEncodeBroadHistoneHuvecH3k09me3A1nRep2.bam.bai |
| HUVEC | H3K9me3 | GSM1003517 | wgEncodeBroadHistoneHuvecH3k09me3A1nRep2.bam |
| HUVEC | Input | GSM733715 | wgEncodeBroadHistoneHuvecControlStdA1nRep1.bam.bai |
| HUVEC | Input | GSM733715 | wgEncodeBroadHistoneHuvecControlStdA1nRep1.bam |
| HUVEC | Input | GSM733715 | wgEncodeBroadHistoneHuvecControlStdA1nRep2.bam.bai |
| HUVEC | Input | GSM733715 | wgEncodeBroadHistoneHuvecControlStdA1nRep2.bam |
| HUVEC | Input | GSM733715 | wgEncodeBroadHistoneHuvecControlStdA1nRep3.bam.bai |
| HUVEC | Input | GSM733715 | wgEncodeBroadHistoneHuvecControlStdA1nRep3.bam |
| NHEK | H3K27ac | GSM733674 | wgEncodeBroadHistoneNhekH3k27acStdA1nRep1.bam.bai |
| NHEK | H3K27ac | GSM733674 | wgEncodeBroadHistoneNhekH3k27acStdA1nRep1.bam |
| NHEK | H3K27ac | GSM733674 | wgEncodeBroadHistoneNhekH3k27acStdA1nRep2.bam.bai |
| NHEK | H3K27ac | GSM733674 | wgEncodeBroadHistoneNhekH3k27acStdA1nRep2.bam |
| NHEK | H3K27ac | GSM733674 | wgEncodeBroadHistoneNhekH3k27acStdA1nRep3.bam.bai |
| NHEK | H3K27ac | GSM733674 | wgEncodeBroadHistoneNhekH3k27acStdA1nRep3.bam |
| NHEK | H3K27me3 | GSM733701 | wgEncodeBroadHistoneNhekH3k27me3StdA1nRep1.bam.bai |
| NHEK | H3K27me3 | GSM733701 | wgEncodeBroadHistoneNhekH3k27me3StdA1nRep1.bam |
| NHEK | H3K27me3 | GSM733701 | wgEncodeBroadHistoneNhekH3k27me3StdA1nRep2.bam.bai |
| NHEK | H3K27me3 | GSM733701 | wgEncodeBroadHistoneNhekH3k27me3StdA1nRep2.bam |
| NHEK | H3K27me3 | GSM733701 | wgEncodeBroadHistoneNhekH3k27me3StdA1nRep3.bam.bai |
| NHEK | H3K27me3 | GSM733701 | wgEncodeBroadHistoneNhekH3k27me3StdA1nRep3.bam |
| NHEK | H3K36me3 | GSM733726 | wgEncodeBroadHistoneNhekH3k36me3StdA1nRep1.bam.bai |
| NHEK | H3K36me3 | GSM733726 | wgEncodeBroadHistoneNhekH3k36me3StdA1nRep1.bam |
| NHEK | H3K36me3 | GSM733726 | wgEncodeBroadHistoneNhekH3k36me3StdA1nRep2.bam.bai |
| NHEK | H3K36me3 | GSM733726 | wgEncodeBroadHistoneNhekH3k36me3StdA1nRep2.bam |
| NHEK | H3K36me3 | GSM733726 | wgEncodeBroadHistoneNhekH3k36me3StdA1nRep3.bam.bai |
| NHEK | H3K36me3 | GSM733726 | wgEncodeBroadHistoneNhekH3k36me3StdA1nRep3.bam |
| NHEK | H3K4me1 | GSM733698 | wgEncodeBroadHistoneNhekH3k4me1StdA1nRep1.bam.bai |
| NHEK | H3K4me1 | GSM733698 | wgEncodeBroadHistoneNhekH3k4me1StdA1nRep1.bam |
| NHEK | H3K4me1 | GSM733698 | wgEncodeBroadHistoneNhekH3k4me1StdA1nRep2.bam.bai |
| NHEK | H3K4me1 | GSM733698 | wgEncodeBroadHistoneNhekH3k4me1StdA1nRep2.bam |

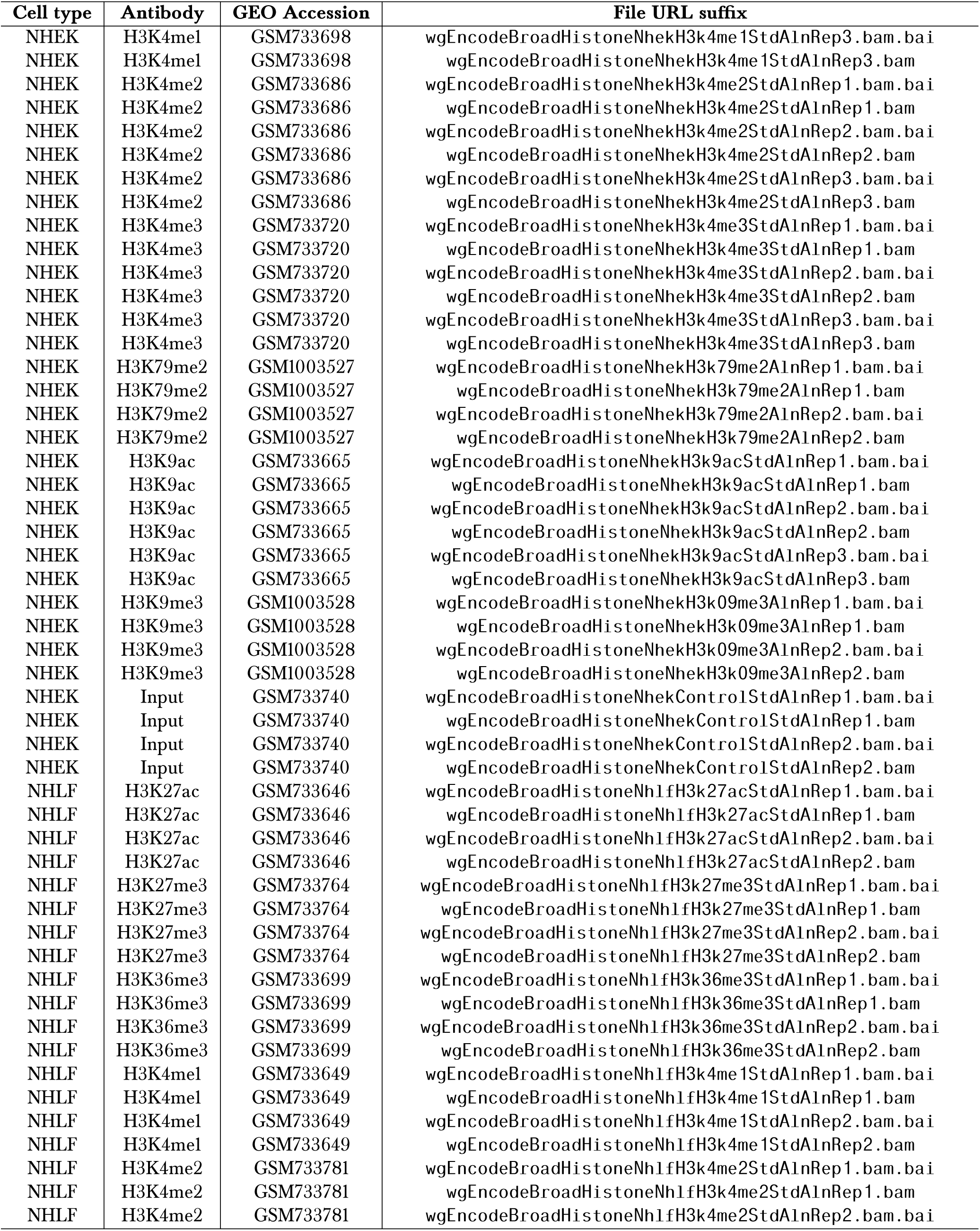

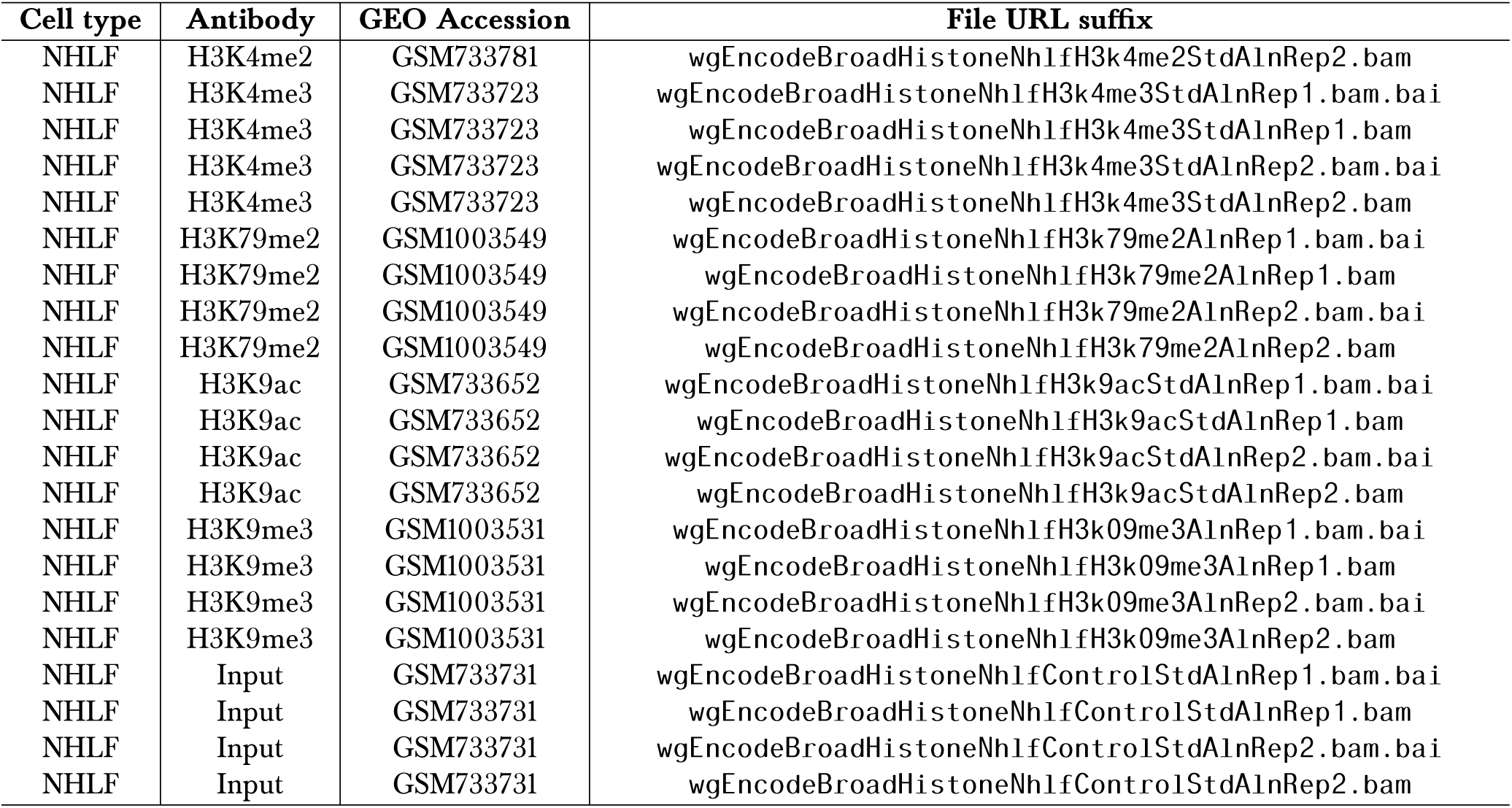

#### *Homo sapiens* source data of RNA-seq transcript abundance in FPKM (GTF files) [54]

For downloading, the URL must be constructed by adding the following prefix to each file listed: ftp://hgdownload.cse.ucsc.edu/goldenPath/hg19/encodeDCC/wgEncodeCaltechRnaSeq/

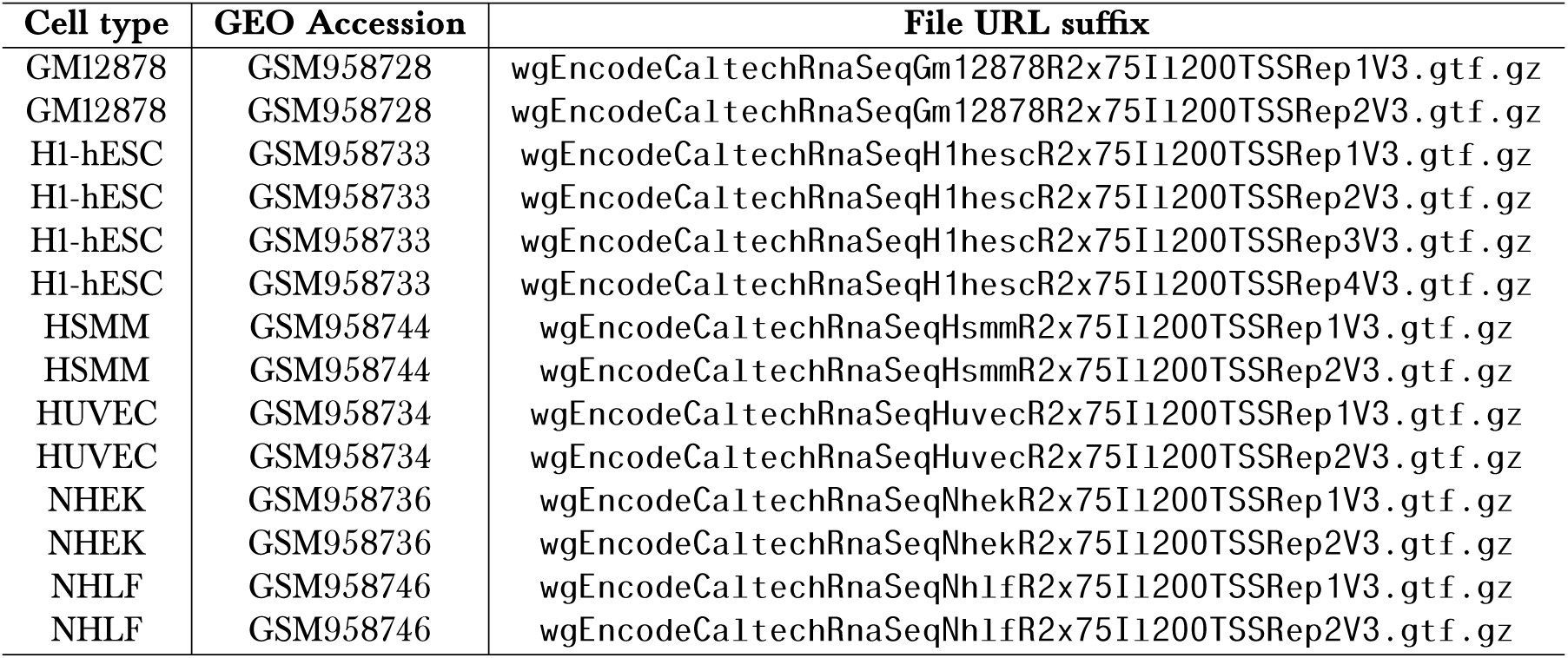

#### *Mus musculus* source data of ChIP-seq on histone H3 modifications (SRA files) [55, 53]

For downloading, the URL must be constructed by adding the following prefix to each file listed:

~~~
ftp://ftp-trace.ncbi.nlm.nih.gov/sra/sra-instant/reads/ByRun/sra/SRR/
~~~

| Cell type | Antibody | Rep # | GEO Accession | File URL suffix |
| --- | --- | --- | --- | --- |
| E14 | IgG | 1 | GSM881345 | SRR414/SRR414932/SRR414932.sra |
| E14-day0 | H3K27ac | 1 | GSM881349 | SRR414/SRR414936/SRR414936.sra |
| E14-day0 | H3K27me3 | 1 | GSM881350 | SRR414/SRR414937/SRR414937.sra |
| E14-day0 | H3K36me3 | 1 | GSM881351 | SRR414/SRR414938/SRR414938.sra |
| E14-day0 | H3K4me1 | 1 | GSM881352 | SRR414/SRR414939/SRR414939.sra |
| E14-day0 | H3K4me3 | 1 | GSM881354 | SRR414/SRR414941/SRR414941.sra |
| E14-day4 | H3K27ac | 1 | GSM881357 | SRR414/SRR414945/SRR414945.sra |
| E14-day4 | H3K27me3 | 1 | GSM881358 | SRR414/SRR414946/SRR414946.sra |
| E14-day4 | H3K36me3 | 1 | GSM881359 | SRR414/SRR414947/SRR414947.sra |
| E14-day4 | H3K4me1 | 1 | GSM881360 | SRR414/SRR414948/SRR414948.sra |
| E14-day4 | H3K4me3 | 1 | GSM881362 | SRR414/SRR414950/SRR414950.sra |
| E14-day6 | H3K27ac | 1 | GSM881366 | SRR414/SRR414955/SRR414955.sra |
| E14-day6 | H3K27me3 | 1 | GSM881367 | SRR414/SRR414956/SRR414956.sra |
| E14-day6 | H3K36me3 | 1 | GSM881368 | SRR414/SRR414957/SRR414957.sra |
| E14-day6 | H3K4me1 | 1 | GSM881369 | SRR414/SRR414958/SRR414958.sra |
| E14-day6 | H3K4me3 | 1 | GSM881371 | SRR414/SRR414960/SRR414960.sra |
| Heart (8 wks/o) | H3K27ac | 1 | GSM1000093 | SRR566/SRR566827/SRR566827.sra |
| Heart (8 wks/o) | H3K27ac | 2 | GSM1000093 | SRR566/SRR566828/SRR566828.sra |
| Heart (8 wks/o) | H3K27me3 | 1 | GSM1000131 | SRR566/SRR566903/SRR566903.sra |
| Heart (8 wks/o) | H3K27me3 | 2 | GSM1000131 | SRR566/SRR566904/SRR566904.sra |
| Heart (8 wks/o) | H3K36me3 | 1 | GSM1000130 | SRR566/SRR566901/SRR566901.sra |
| Heart (8 wks/o) | H3K36me3 | 2 | GSM1000130 | SRR566/SRR566902/SRR566902.sra |
| Heart (8 wks/o) | H3K4me1 | 1 | GSM769025 | SRR317/SRR317255/SRR317255.sra |
| Heart (8 wks/o) | H3K4me1 | 2 | GSM769025 | SRR317/SRR317256/SRR317256.sra |
| Heart (8 wks/o) | H3K4me3 | 1 | GSM769017 | SRR317/SRR317239/SRR317239.sra |
| Heart (8 wks/o) | H3K4me3 | 2 | GSM769017 | SRR317/SRR317240/SRR317240.sra |
| Heart (8 wks/o) | Input | 1 | GSM769032 | SRR317/SRR317269/SRR317269.sra |
| Heart (8 wks/o) | Input | 2 | GSM769032 | SRR317/SRR317270/SRR317270.sra |
| Liver (8 wks/o) | H3K27ac | 1 | GSM1000140 | SRR566/SRR566921/SRR566921.sra |
| Liver (8 wks/o) | H3K27ac | 2 | GSM1000140 | SRR566/SRR566922/SRR566922.sra |
| Liver (8 wks/o) | H3K27me3 | 1 | GSM1000150 | SRR566/SRR566941/SRR566941.sra |
| Liver (8 wks/o) | H3K27me3 | 2 | GSM1000150 | SRR566/SRR566942/SRR566942.sra |
| Liver (8 wks/o) | H3K36me3 | 1 | GSM1000151 | SRR566/SRR566943/SRR566943.sra |
| Liver (8 wks/o) | H3K36me3 | 2 | GSM1000151 | SRR566/SRR566944/SRR566944.sra |
| Liver (8 wks/o) | H3K4me1 | 1 | GSM769015 | SRR317/SRR317235/SRR317235.sra |
| Liver (8 wks/o) | H3K4me1 | 2 | GSM769015 | SRR317/SRR317236/SRR317236.sra |
| Liver (8 wks/o) | H3K4me3 | 1 | GSM769014 | SRR317/SRR317233/SRR317233.sra |
| Liver (8 wks/o) | H3K4me3 | 2 | GSM769014 | SRR317/SRR317234/SRR317234.sra |
| Liver (8 wks/o) | Input | 1 | GSM769034 | SRR317/SRR317273/SRR317273.sra |
| Liver (8 wks/o) | Input | 2 | GSM769034 | SRR317/SRR317274/SRR317274.sra |

#### *Mus musculus* source data of RNA-seq (BAM files) [55, 53]

For downloading, the URL must be constructed by adding one of the two following prefixes to each file listed:

~~~
1. ftp://ftp.ncbi.nlm.nih.gov/geo/samples/GSM881nnn/
2. ftp://hgdownload.cse.ucsc.edu/goldenPath/mm9/encodeDCC/wgEncodeLicrRnaSeq/
~~~

| Cell type | Rep # | GEO Accession | File URL suffix |
| --- | --- | --- | --- |
| E14-day0 | 1 | GSM881355 | [*prefix_1*]GSM881355/suppl/GSM881355_E14_RNA.bam.gz |
| E14-day4 | 1 | GSM881364 | [*prefix_1*]GSM881364/suppl/GSM881364_E14_RNA_d4.bam.gz |
| E14-day6 | 1 | GSM881373 | [*prefix_1*]GSM881373/suppl/GSM881373_E14_RNA_d6.bam.gz |
| Heart (8 wks/o) | 1 | GSM929707 | [*prefix_2*]wgEncodeLicrRnaSeqHeartCellPapMAdu1t8wksC57b16A1nRep1.bam |
| Heart (8 wks/o) | 2 | GSM929707 | [*prefix_2*]wgEncodeLicrRnaSeqHeartCellPapMAdu1t8wksC57b16A1nRep2.bam |
| Liver (8 wks/o) | 1 | GSM929711 | [*prefix_2*]wgEncodeLicrRnaSeqLiverCellPapMAdu1t8wksC57b16A1nRep1.bam |
| Liver (8 wks/o) | 2 | GSM929711 | [*prefix_2*]wgEncodeLicrRnaSeqLiverCellPapMAdu1t8wksC57b16A1nRep2.bam |

#### *Drosophila melanogaster* source data of ChIP-seq on histone H3 modifications (SRA files) [49, 51]

For downloading, the URL must be constructed by adding the following prefix to each file listed:

~~~
ftp://ftp-trace.ncbi.nlm.nih.gov/sra/sra-instant/reads/ByRun/sra/SRR/SRR030/
~~~

| Developmental time point/period | Antibody | GEO Accession | File URL suffix |
| --- | --- | --- | --- |
| 0-4h embryos | H3K27ac | GSM401407 | SRR030295/SRR030295.sra |
| 0-4h embryos | H3K27me3 | GSM439448 | SRR030360/SRR030360.sra |
| 0-4h embryos | H3K4me1 | GSM401409 | SRR030297/SRR030297.sra |
| 0-4h embryos | H3K4me3 | GSM400656 | SRR030269/SRR030269.sra |
| 0-4h embryos | H3K9ac | GSM401408 | SRR030296/SRR030296.sra |
| 0-4h embryos | H3K9me3 | GSM439457 | SRR030369/SRR030369.sra |
| 0-4h embryos | Input | GSM400657 | SRR030270/SRR030270.sra |
| 4-8h embryos | H3K27ac | GSM401404 | SRR030292/SRR030292.sra |
| 4-8h embryos | H3K27me3 | GSM439447 | SRR030359/SRR030359.sra |
| 4-8h embryos | H3K4me1 | GSM401406 | SRR030294/SRR030294.sra |
| 4-8h embryos | H3K4me3 | GSM400674 | SRR030287/SRR030287.sra |
| 4-8h embryos | H3K9ac | GSM401405 | SRR030293/SRR030293.sra |
| 4-8h embryos | H3K9me3 | GSM439456 | SRR030368/SRR030368.sra |
| 4-8h embryos | Input | GSM400675 | SRR030288/SRR030288.sra |
| 8-12h embryos | H3K27ac | GSM432583 | SRR030332/SRR030332.sra |
| 8-12h embryos | H3K27me3 | GSM439446 | SRR030358/SRR030358.sra |
| 8-12h embryos | H3K4me1 | GSM432593 | SRR030342/SRR030342.sra |
| 8-12h embryos | H3K4me3 | GSM432585 | SRR030334/SRR030334.sra |
| 8-12h embryos | H3K9ac | GSM432592 | SRR030341/SRR030341.sra |
| 8-12h embryos | H3K9me3 | GSM439455 | SRR030367/SRR030367.sra |
| 8-12h embryos | Input | GSM432636 | SRR030346/SRR030346.sra |
| 12-16h embryos | H3K27ac | GSM432582 | SRR030331/SRR030331.sra |
| 12-16h embryos | H3K27me3 | GSM439445 | SRR030357/SRR030357.sra |
| 12-16h embryos | H3K4me1 | GSM432591 | SRR030340/SRR030340.sra |
| 12-16h embryos | H3K4me3 | GSM432580 | SRR030329/SRR030329.sra |

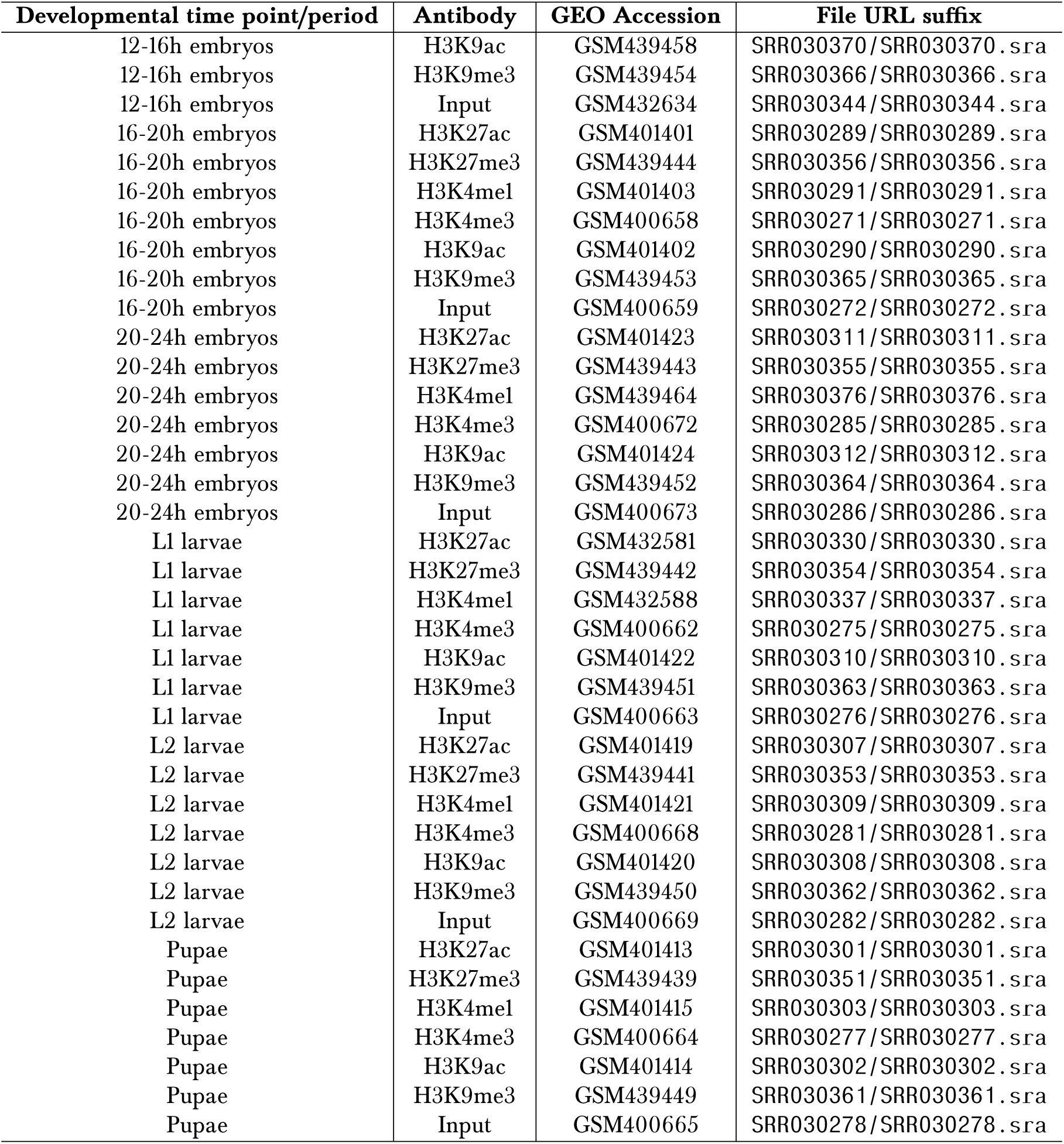

| Developmental time point/period | Antibody | GEO Accession | File URL suffix |
| --- | --- | --- | --- |
| 12-16h embryos | H3K9ac | GSM439458 | SRR030370/SRR030370.sra |
| 12-16h embryos | H3K9me3 | GSM439454 | SRR030366/SRR030366.sra |
| 12-16h embryos | Input | GSM432634 | SRR030344/SRR030344.sra |
| 16-20h embryos | H3K27ac | GSM401401 | SRR030289/SRR030289.sra |
| 16-20h embryos | H3K27me3 | GSM439444 | SRR030356/SRR030356.sra |
| 16-20h embryos | H3K4me1 | GSM401403 | SRR030291/SRR030291.sra |
| 16-20h embryos | H3K4me3 | GSM400658 | SRR030271/SRR030271.sra |
| 16-20h embryos | H3K9ac | GSM401402 | SRR030290/SRR030290.sra |
| 16-20h embryos | H3K9me3 | GSM439453 | SRR030365/SRR030365.sra |
| 16-20h embryos | Input | GSM400659 | SRR030272/SRR030272.sra |
| 20-24h embryos | H3K27ac | GSM401423 | SRR030311/SRR030311.sra |
| 20-24h embryos | H3K27me3 | GSM439443 | SRR030355/SRR030355.sra |
| 20-24h embryos | H3K4me1 | GSM439464 | SRR030376/SRR030376.sra |
| 20-24h embryos | H3K4me3 | GSM400672 | SRR030285/SRR030285.sra |
| 20-24h embryos | H3K9ac | GSM401424 | SRR030312/SRR030312.sra |
| 20-24h embryos | H3K9me3 | GSM439452 | SRR030364/SRR030364.sra |
| 20-24h embryos | Input | GSM400673 | SRR030286/SRR030286.sra |
| L1 larvae | H3K27ac | GSM432581 | SRR030330/SRR030330.sra |
| L1 larvae | H3K27me3 | GSM439442 | SRR030354/SRR030354.sra |
| L1 larvae | H3K4me1 | GSM432588 | SRR030337/SRR030337.sra |
| L1 larvae | H3K4me3 | GSM400662 | SRR030275/SRR030275.sra |
| L1 larvae | H3K9ac | GSM401422 | SRR030310/SRR030310.sra |
| L1 larvae | H3K9me3 | GSM439451 | SRR030363/SRR030363.sra |
| L1 larvae | Input | GSM400663 | SRR030276/SRR030276.sra |
| L2 larvae | H3K27ac | GSM401419 | SRR030307/SRR030307.sra |
| L2 larvae | H3K27me3 | GSM439441 | SRR030353/SRR030353.sra |
| L2 larvae | H3K4me1 | GSM401421 | SRR030309/SRR030309.sra |
| L2 larvae | H3K4me3 | GSM400668 | SRR030281/SRR030281.sra |
| L2 larvae | H3K9ac | GSM401420 | SRR030308/SRR030308.sra |
| L2 larvae | H3K9me3 | GSM439450 | SRR030362/SRR030362.sra |
| L2 larvae | Input | GSM400669 | SRR030282/SRR030282.sra |
| Pupae | H3K27ac | GSM401413 | SRR030301/SRR030301.sra |
| Pupae | H3K27me3 | GSM439439 | SRR030351/SRR030351.sra |
| Pupae | H3K4me1 | GSM401415 | SRR030303/SRR030303.sra |
| Pupae | H3K4me3 | GSM400664 | SRR030277/SRR030277.sra |
| Pupae | H3K9ac | GSM401414 | SRR030302/SRR030302.sra |
| Pupae | H3K9me3 | GSM439449 | SRR030361/SRR030361.sra |
| Pupae | Input | GSM400665 | SRR030278/SRR030278.sra |

#### *Drosophila melanogaster* source data of RNA-seq (SAM files) [49, 51]

For downloading, the URL must be constructed by adding the following prefix to each file listed:

~~~
ftp://data.modencode.org/all_files/dmel-signal-1/
~~~

| Developmental time point/period | GEO Accession | File URL suffix |
| --- | --- | --- |
| 0-4h embryos | GSM451806 | 2010_0-4_accepted_hits.sam.gz |
| 4-8h embryos | GSM451809 | 2019_4-8_accepted_hits.sam.gz |
| 8-12h embryos | GSM451808 | 2020_8-12_accepted_hits.sam.gz |
| 12-16h embryos | GSM451803 | 2021_12-16_accepted_hits.sam.gz |
| 16-20h embryos | GSM451807 | 2022_16-20_accepted_hits.sam.gz |
| 20-24h embryos | GSM451810 | 2023_20-24_accepted_hits.sam.gz |
| L1 larvae | GSM451811 | 2024_L1_accepted_hits.sam.gz |
| L2 larvae | GSM453867 | 2025_L2_accepted_hits.sam.gz |
| Pupae | GSM451813 | 2030_Pupae_accepted_hits.sam.gz |

